# ROS production by localized SCHENGEN receptor module drives lignification at subcellular precision

**DOI:** 10.1101/818997

**Authors:** Satoshi Fujita, Damien De Bellis, Kai H. Edel, Phillipp Köster, Tonni Grube Andersen, Emanuel Schmid-Siegert, Valérie Dénervaud Tendon, Alexandre Pfister, Peter Marhavý, Robertas Ursache, Verónica G. Doblas, Jean Daraspe, Audrey Creff, Gwyneth Ingram, Jörg Kudla, Niko Geldner

**Author notes:** Correspondence to (S.F.) and (N.G.).

## Abstract

Production of reactive-oxygen species (ROS) by NADPH oxidases (NOXs) impacts many processes in animals and plants and many plant receptor pathways involve rapid, NOX-dependent increases of ROS. Yet, their general reactivity has made it challenging to pinpoint the precise role and direct cellular targets of ROS. A well-understood ROS target in plants are lignin peroxidases in the cell wall. Lignin can be deposited with exquisite spatial control, but the underlying mechanisms have remained elusive. Here we establish a full kinase signaling relay that exerts direct, spatial control over ROS production and lignification within the cell wall. We show that polar localization of a single kinase component is crucial for pathway function. Our data indicates that an intersection of more broadly localized components allows for micrometer-scale precision of lignification and that this system is triggered through initiation of ROS production as a critical peroxidase co-substrate.

---

As in animals, NADPH oxidase-produced ROS in plants is important for a multitude of processes and the number of NADPH oxidase genes (10 in Arabidopsis, called *RESPIRATORY BURST OXIDASE HOMOLOGs, RBOHs, A-J*) suggests a high complexity of regulation of ROS production in plants. Among its many roles, ROS-dependent regulation of plant cell wall structure and function is considered to be among its most critical ^1^. The cell wall is the nano-structured, sugar-based, pressure-resisting extracellular matrix of plants and NOXs are thought to be the predominant ROS source in this compartment (also termed apoplast) ^1^.

A staggering number of kinases have been shown to regulate plant NOXs and the activation mechanism of NOX-dependent ROS production is well established, especially in response to microbial pattern-recognition by immune receptors ^2^. However, the specific role and direct molecular targets of ROS during microbial pattern-recognition have remained elusive ^3^. The same applies for the central role of ROS in tip growing cells, such as root hairs or pollen tubes, where ROS is thought to be part of an intricate oscillation of cell wall stiffening and loosening, aimed at allowing cell wall expansion without catastrophic collapse ^4,5^. In this case, ROS is proposed to be important for counteracting cell wall loosening pH decreases, but it is again unclear what direct targets of ROS would mediate cell wall stiffening. Cell wall lignification by apoplastic peroxidases, can therefore be considered as the most well-established role of ROS, where the peroxidases themselves are the direct “ROS targets”, using it as a co-substrate for the oxidation of mono-lignols ^6,7^. In the case of lignification, however, a molecularly-defined signaling pathway that induces ROS production during lignification has not been defined. A few years ago, our group identified a specific NADPH oxidase, RBOHF, to be required for the localized formation of lignin in the root endodermis ^8^. Lignin is a poly-phenolic polymer that is generated by the radical-coupling of mono-lignols, oxidized through the action of ROS-dependent peroxidases, as well as laccases ^7^. The hydrophobic lignin polymer impregnates the cellulosic cell wall of plants, rendering it unextendible and highly resistant to degradation. Lignin in the root endodermis is deposited in a central, longitudinal band around every endodermal cell. Named Casparian strips (CS), these ring-like lignin structures fuse into a supracellular network, establishing a tissue-wide, extracellular diffusion barrier (Fig. 1A), analogous to epithelial tight junctions in animals ^9,10^. Functionality of this barrier can be easily visualized by a block of penetration of a fluorescent cell wall dye, propidium iodide, into the vasculature ^11,12^. CS localization occurs through the action of CASPARIAN STRIP DOMAIN PROTEINS (CASPs), 4-TM proteins, which form a highly scaffolded transmembrane protein platform, assembling RBOHF and cell wall peroxidases and other proteins at the Casparian strip domain (CSD) ^8,13,14^. CSD formation and lignin deposition is coordinated such that the aligned rings of endodermal neighbors’ fuse, leading to a supracellular network that seals the extracellular space between endodermal cells, generating a tissue-wide diffusion barrier.

**Fig. 1.**
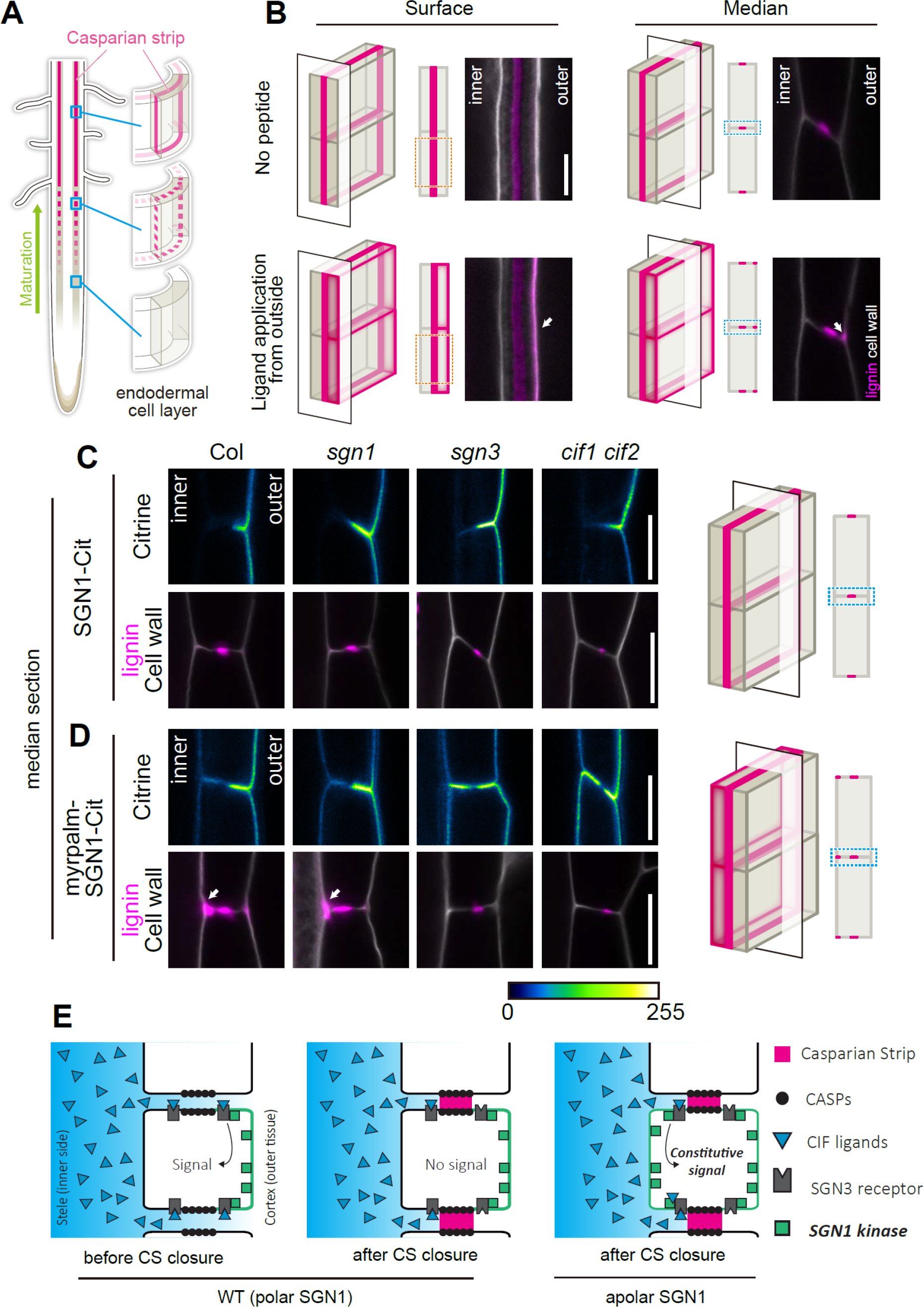
Apolar SGN1 leads to ectopic lignin accumulation in endodermal cells. (A) Schematic of Casparian strip development (magenta). Casparian strips start to appear as centrally aligned discontinuous dots in the endodermal cell layer, progressing into a network of fused rings functioning as a root apoplastic barrier. (B) Lignin accumulation patterns at endodermal surface or median positions with or without the CIF2 (Casparian strip integrity factor 2) ligand. Lignin and cellulosic (unmodified) cell walls are stained with Basic Fuchsin and Calcofluor White, shown in magenta and white respectively. Schematics are indicating the position of optical sections in a 3D illustration. Scale Bar = 5 *μ* m. (C,D) Localization of SGN1-Cit and lignin deposition patterns in *pCASP1∷SGN1-Cit* lines in wild-type (Col) and different mutant backgrounds (*sgn1, sgn3, cif1 cif2*) (C). myrpalm-SGN1-Cit localization and lignin deposition patterns in *pCASP1∷myrpalm-SGN1-Cit* lines (D). Lignin (Basic Fuchsin) and cell walls (Calcoflour White) are shown in magenta and white respectively. Schematics are indicating the position of optical sections in a 3D illustration. Scale bars = 5 *μ* m. (E) Schematic illustrating how signal activation can be governed by SGN1 localization and peptide ligand diffusion from the stele. “inner” designates the stele-facing endodermal surface, “outer”, the cortex-facing surface.

More recently, we identified a pair of peptide ligands, a Leucine-rich repeat receptor-like kinase (LRR-RLK) and a cytoplasmic kinase, whose phenotypes, genetic interaction and specific subcellular localizations led us to propose that they combine into a barrier surveillance pathway. Previous reports had shown that the CIF1/2 (CASPARIAN STRIP INTEGRITY FACTORs 1/2) peptides are SCHENGEN3 (SGN3) (also called GASSHO1 (GSO1)) ligands and that the SGN1 and SGN3 kinases govern Casparian strip integrity ^15–18^. *cif1 cif2* and *sgn1, sgn3* mutants have similar, discontinuous CS, caused by a discontinuous CSD, as well as a conspicuous absence, or strong attenuation of, compensatory lignification and suberization observed in other CS mutants ^13,15,18–21^. Their phenotypic similarities suggested that these factors act in one pathway. CIF1 and 2 peptides do not express in the endodermis, where the CS is formed, but in the stele. In contrast, both SGN1 and SGN3 present specific localization on the endodermal plasma membrane; SGN3 receptor-like kinase resides along both sides of the CS, while palmitoylated SGN1 polarly localizes on cortex facing (outer) plasma membranes ^15^. Remarkably, their localization overlaps only in a small region next to the cortex-facing side of the CS ^16^. This would require peptides from the stele to diffuse across the CS in order to access the signaling complex. This is only possible while the CS is still permeable ^15^ (Fig. 1E).

This pathway would therefore allow the endodermal cell layer to probe the tissue-wide integrity of its extracellular diffusion barrier and to respond to barrier defects through compensatory over-lignification ^15^. Here we demonstrate that the receptor, cytoplasmic kinase and NADPH oxidase molecularly connect into one pathway, with great resemblance to plant immune signaling pathways, but whose direct action is to locally produce ROS for localized lignification. We establish the crucial importance of the restricted subcellular localization of its components and demonstrate that stimulation of this signaling pathway additionally leads to strong transcriptional activation of target genes that further drive and sustain endodermal lignification, as well as suberization and endodermal sub-domain formation and differentiation. We thus provide a full molecular circuitry in which an endogenous peptide from the stele stimulates localized signaling kinases and NADPH oxidases in the endodermis, causing extracellular ROS production at micrometer-scale precision and a precisely localized lignification of the plant cell wall.

## RESULTS

We generated a new *cif1-2 cif2-2* double mutant allele by CRISPR-Cas9 in a pure Col background, because the previous *cif1-1 cif2-1* double mutant allele ^17^ was a mixture of Ws and Col alleles. The new CRISPR allele was complemented by CIF1 or CIF2 application, visualized by PI (propidium iodide) uptake assays or reconstitution of CASP1 membrane domain connectivity (Fig. S1D,E and Fig. S2A).

### Apolar SGN1 kinase leads to constitutive barrier defect signaling

Central to the barrier surveillance model is the polar localization of SGN1, which is thought to limit the potential for signal activation to the cortex side of the endodermis, requiring passage of CIF peptides across the CS region (Fig. 1A,E). Consistently, application of peptide ligand to the media, leading to stimulation from the outside, causes overlignification at the cortex-facing endodermal edges (Fig. 1B). In an attempt to falsify the model we had proposed and to interrogate the importance of SGN1 polar localization, we generated a SGN1 variant that localized in an apolar fashion, by adding a myristoylation (myr) and palmitoylation (palm) motifs on the N terminus ^22^. This myrpalm SGN1-mCitrine (Cit) variant was expressed under the control of the endodermis-specific CASP1 promoter, which is strongly active during Casparian strip formation and complemented the *sgn1* barrier phenotype (Fig. S1A). *In planta*, the wild-type SGN1-Cit variant resides polarly on the cortex-facing side of endodermal cells, while myrpalm SGN1-Cit was observed at both sides of the endodermal plasma membranes, even though preferentially accumulation at the cortex side could still be observed (Fig. 1C, 1D). Both variants were excluded from the central position where the Casparian strip domain is formed (Fig. S1B) and the localization patterns of the two variants did not change when introgressed into *sgn1*, *sgn3* or *cif1 cif2* mutants (Fig. 1C, 1D). An apolar SGN1 localization would allow SGN3 to encounter SGN1 also on the stele-facing side, not only on the cortex side (Fig. 1E). This should lead to constitutive signal activation in the absence of barrier defects, because the CIF peptides would now be able to access a SGN3/SGN1 signaling module on the stele-facing side without crossing the barrier. Indeed, we found that the apolar SGN1 variant caused both, ectopic lignin deposition and precocious suberization in endodermal cells (Fig. 1D, S1F, G), as previously described for endodermal barrier mutants. Yet, no barrier defect was observed in the lines complemented with apolar SGN1 (Fig. S1A) and, consistently, we found CASP1-mCherry distribution to be normal in these lines, forming a continuous band in the central position of the endodermal cells, indistinguishable from wild-type (Fig. S1B). This indicates that presence of SGN1 at the plasma membrane to the inside of the CS leads to signaling in absence of barrier defects. CASP1-driven, wild type SGN1 lines as a control, complemented the mutant (Fig. S1A) and did not cause any changes in lignin accumulation pattern (Fig. 1C). Interestingly, the apolar SGN1 lines accumulated lignin mainly on the stele-facing edges of the endodermal cell walls, as opposed to the cortical lignin deposition observed by ectopic ligand treatment (compare Fig. 1B with Fig. 1C, arrows). The ectopic lignin deposition in apolar SGN1 lines was fully dependent on the presence of receptor and ligand, as both in *sgn3* and *cif1 cif2* mutants, no excess lignification at the stele-facing side could be observed in apolar SGN1 lines (Fig. 1D). This strongly suggests that mislocalized SGN1 does not become constitutively active, but leads to continuous, ectopic transduction of CIF1/2 signals through SGN3. An apolar, but kinase-dead variant of SGN1 was also unable to induce ectopic lignification (Fig. S1C), suggesting that a phosphorylation relay downstream of SGN1 is necessary for lignification.

### SGN1 is a downstream component of the CIF/SGN3 pathway

Previous data showed that *sgn1* is less sensitive to high doses of externally applied CIF peptide, consistent with a role of SGN1 downstream of the SGN3 receptor ^15^, at least with respect to CIF-induced excess lignification. In order to address whether SGN1 is indeed a generally required downstream component of the CIF/SGN3 pathway during CS formation, we evaluated whether the *sgn1* mutant is resistant to complementation by CIF2 peptide treatment. In contrast to the full complementation of the discontinuous CS domain of *cif1 cif2* double mutants, the *sgn1* mutant did not show full restoration of domain integrity, neither on 10 nM or 100 nM CIF2 medium (Fig. 2A, S2A). While some degree of rescue occurred, only about 50% of discontinuities were rescued. This was corroborated by testing CS functionality using the propidium iodide (PI) assay. Only a weak complementation of barrier formation was observed even when grown on 100 nM CIF2 medium (Fig. 2B). Our results indicate that SGN1 functions downstream of CIFs/SGN3, but suggest that additional factors can partially compensate for its absence, most probably homologs of the extended RLCKVII family to which SGN1 belongs.

**Fig. 2.**
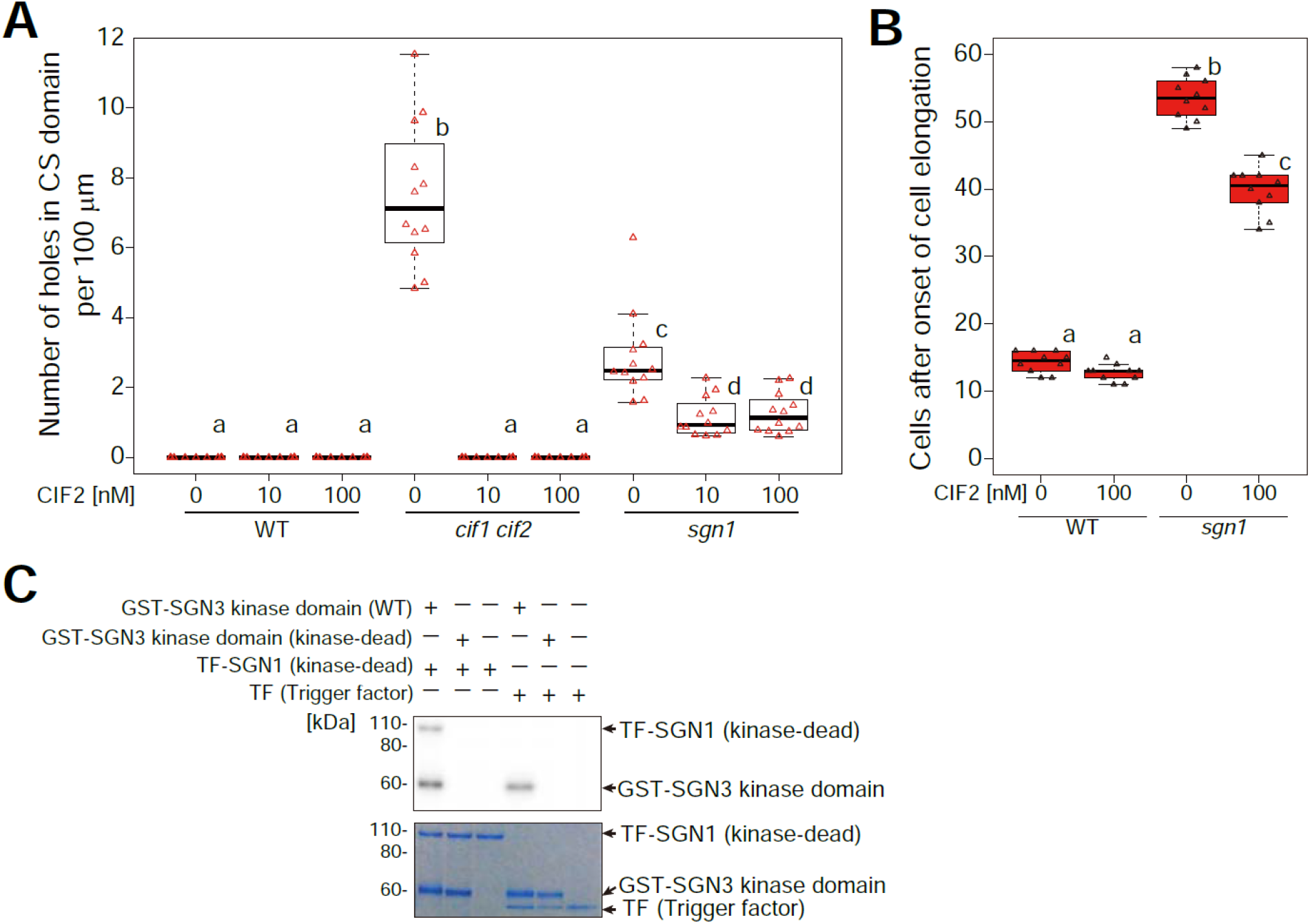
SGN1 acts as a transducer of CIF2 signaling and is phosphorylated by the SGN3 receptor. (A) Quantification of defects in CSD formation as number of holes per 100 *μ* m in CASP1-GFP domain at around 10 cells after onset of CASP1-GFP expression in 5-day-old seedlings. One-way ANOVA was performed followed by Tukey’s test. Different letters show significant statistical differences. (p<0.05, One-way ANOVA and Tukey’s test, n= 12 from each condition). (B) Propidium iodide (PI) penetration assay in the presence or the absence of CIF2. CS barrier function was scored as exclusion of PI signal from the inner side of endodermal cells. Different letters show significant statistical differences. (p<0.05, One-way ANOVA and Tukey’s test n= 10 from each condition). (C) *In vitro* kinase assay of SGN3 kinase domain against SGN1. Note that a kinase-dead SGN1 variant was used to avoid the auto-phosphorylation activity of SGN1. Representative result of three independent experiments is shown.

We then tested for direct connectivity between SGN3 receptor kinase and SGN1 by carrying out an *in vitro* kinase assay. Some RLCKVII members are reported to be phosphorylated and activated by LRR receptor kinases ^23,24^. A Glutathione S-transferase (GST)-fused SGN3 kinase domain was incubated with a kinase-dead form of a trigger factor (TF)-SGN1 fusion protein - which does not have autophosphorylation activity - in the presence of radioactive ATP. Kinase-dead TF-SGN1 was efficiently phosphorylated by SGN3 kinase domain, but not by a kinase-dead form of SGN3 *in vitro* (Fig. 2C, S2B). This further supports our model that SGN1 is a direct downstream component of the SGN3/CIF pathway.

### Two NADPH oxidases are absolutely required for CIF-induced lignification in the endodermis

Our generation of an apolar SGN1 thus appears to have reconstituted a functional CIF/SGN3 pathway at the stele-facing (inner) endodermal surface, causing ectopic lignification in the absence of barrier defects. Yet, the second intriguing aspect of this manipulation is the observation that lignification occurs almost exclusively at the inner endodermal edges (Fig. 1D), the side where endogenous CIF peptide must be assumed to be present. External treatment, by contrast, leads to predominant lignification at the outer endodermal edges (Fig. 1B). This surprising spatial correlation between the site of signal perception and localized lignification suggests a very direct molecular connection between the two events that would allow to maintain spatial information. In a cell primed for lignification, i.e. with mono-lignol substrates available and polymerizing enzymes expressed, lignification could be simply “switched on” by activating ROS production through NADPH oxidases. Previously, we had found one of the NOXs, RBOHF, to be crucial for lignification at the CS. Intriguingly, RBOHF is the only transmembrane protein known to accumulate at the Casparian strip domain, safe the CASPs themselves ^8^ (Fig. 3A). Moreover, homologous NADPH oxidases, such as RBOHB or RBOHD, also present in the endodermal plasma membrane, are excluded from this domain ^8^ (Fig. 3A). We therefore asked whether CIF peptides would induce lignification through activation of RBOHF as a downstream component. To our surprise, CIF treatment still led to induced lignification in *rbohf*, despite the fact that RBOHF, is strictly required for CS lignification in untreated conditions. Yet, when RBOHD was knocked-out in addition to RBOHF, a complete absence of lignification upon CIF treatment was observed (Fig. 3B). The fact that RBOHD single mutant neither showed defects in developmental CS lignification, nor in CIF-induced lignification, indicates that RBOHF is required for both processes, but that upon strong stimulation with exogenous CIF peptide application, RBOHD can be additionally used. Indeed, using the general NADPH oxidase inhibitor diphenyleneiodonium (DPI) in short-term co-treatment with CIF was also able to fully block CIF-induced ectopic lignification in wild-type with intact Casparian strips, further supporting that NADPH oxidases act downstream in the CIF/SGN3 pathway (Fig. S3A).

**Fig. 3.**
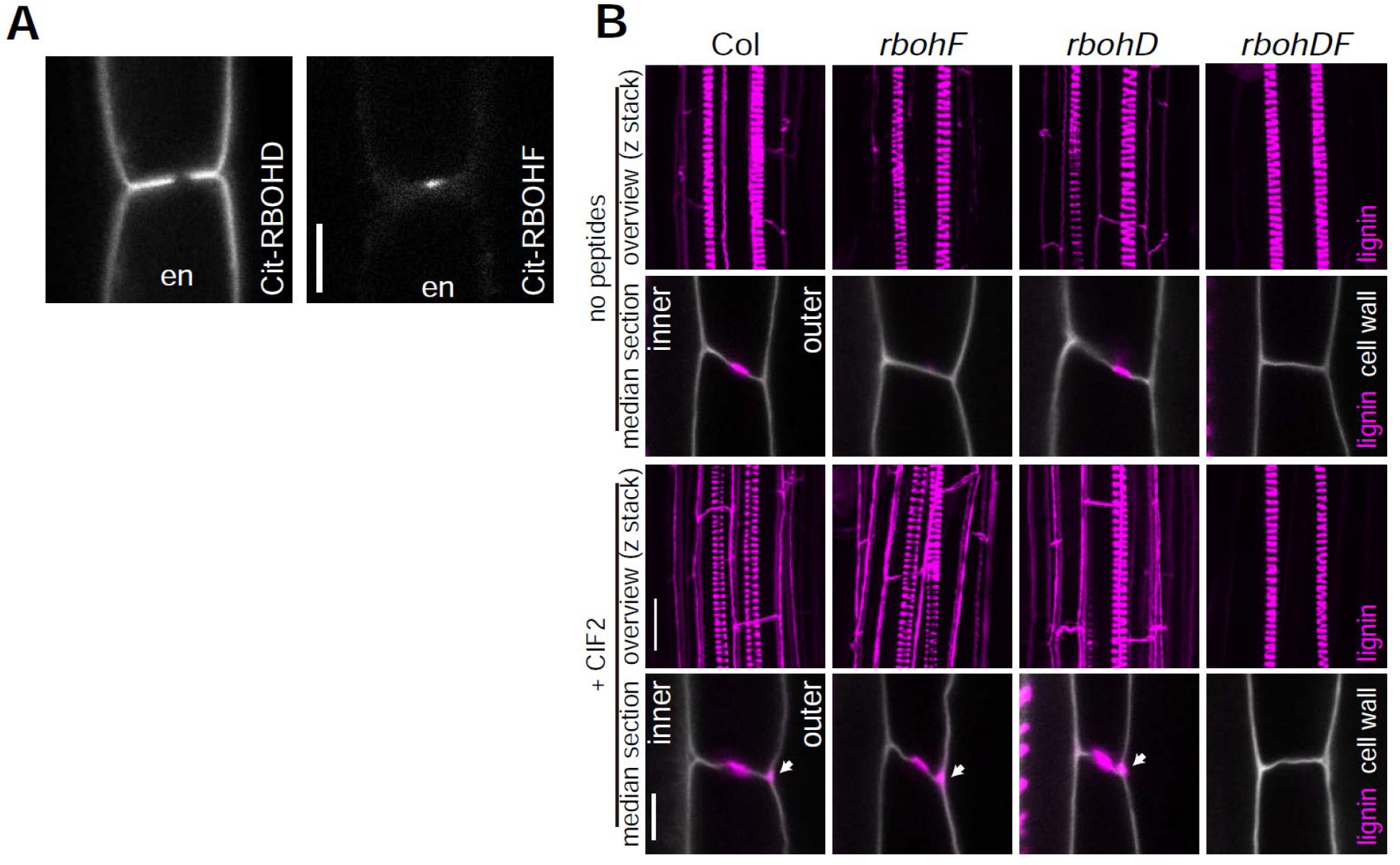
Both RBOHD and F are required for CIF2-induced excess lignin accumulation. (A) Localization of Cit-RBOHD (left) or Cit-RBOHF (right) in endodermal (en) cells. Both proteins were expressed under the control of pCASP1, an endodermal-specific promoter. Scale bar = 5 *μ* m. (B) Lignin accumulation in WT and *rbohD and rbohF* single mutants and a double mutant with or without 2-hour CIF2 peptide treatment. Arrowheads indicate excess lignification. Pictures are shown as overviews (maximum projection) or median sections. Lignin and cell walls are shown with magenta (stained with Basic Fuchsin) and gray (stained with Calcofluor White) respectively. Scale bars = 20 *μ* m (lignin overviews), 5 *μ* m (median sections). “inner” designates the stele-facing endodermal surface, “outer”, the cortex-facing surface.

### CIF2 induces highly localized ROS production through RBOHF and RBOHD

The direct activity of NOX enzymes is not lignification, but production of superoxide (O2^−^) that becomes dismutated to hydrogen peroxide (H_2_O_2_) in the apoplast ^1^. We had previously established that the different subcellular distribution of the SGN3 receptor and the SGN1 kinase only intersect at a very restricted domain at the outer (cortex-facing) edge of the CS ^16^(Fig. 4A). We therefore attempted to visualize whether ROS might be produced locally in response to CIF treatment. Many ways exist to visualize and quantify ROS, but only few allow for high spatial resolution and for discrimination between extracellular and intracellular ROS. An older method, based on ROS-induced cerium precipitation that can be detected using transmission electron microcopy (TEM), has been used by us previously to demonstrate that a highly localized ROS production indeed occurs at the CS and is dependent on NOX activity ^8,25^(Fig. 4B). Using this method, we could detect a strong ROS production in response to CIF2, exclusively at the regions of endodermal-endodermal cell walls outside of the CS, but nowhere along the endodermis-cortex cell walls, which are equally reached by the cerium chloride and where NOX enzymes are also present in the plasma membrane (Fig. 4B,C). Based on this striking spatial coincidence between the SGN3/SGN1 overlap region (Fig. 4A) ^16^ and CIF2-induced ROS production, we developed a procedure to quantitatively assess cerium precipitates and checked whether this localized ROS production is indeed dependent on the SGN3 pathway (Fig. 4D,E, S4A, see also Experimental Procedures section). For SGN3, we found that already steady-state ROS production was undetectable in the mutant and that there was no increase upon CIF2-treatment (Fig. 4D,E). The *sgn1* mutant showed lower, but still detectable steady state ROS levels, but no significant increase upon CIF2-treatment (Fig. 4D,E). Thus, the highly localized ROS accumulation induced by CIF2 is entirely dependent on the localized SGN3/SGN1 receptor module. As in the case of lignification, we observed that CIF2-induced ROS can be produced by either RBOHF or RBOHD, as only the double mutant caused a complete absence of ROS after CIF-stimulation. As expected, but not previously demonstrated, the steady-state ROS production at the CS observed before stimulation was exclusively dependent on RBOHF, but not RBOHD (Fig. 4D,E).

**Fig. 4.**
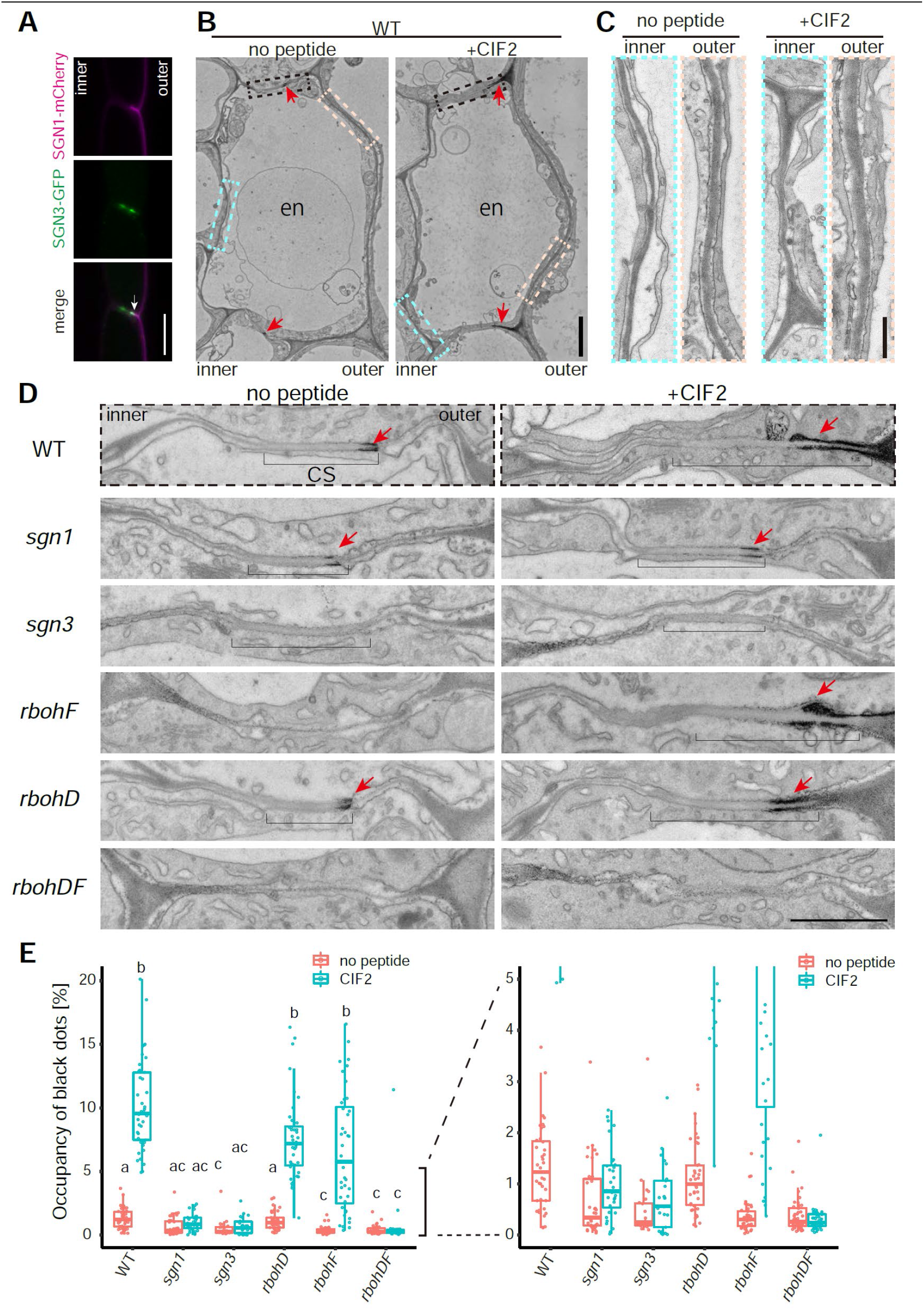
ROS production is enhanced by SGN3/CIFs and requires RBOHD and F. (A) Co-visualization of *pCASP1∷SGN1-mCherry* and *pCASP1∷SGN3-GFP*. Note that their respective localization at the PM has only restricted overlap at the cortical side of the CS domain (Arrow). Scale bar = 10 *μ* m. “inner” designates the stele-facing endodermal surface, “outer”, the cortex-facing surface. (B,C) Overview of endodermal cells after with or without CIF2 treatment. Red arrowheads are indicating ROS production sites and boxes in dotted lines are corresponding to the region in (sky blue and creme boxes in C and black boxes in D). Scale bar = 500 nm. (D) *In situ* H_2_O_2_ detection at Casparian strips in WT, *sgn3*, *sgn1*, *rbohF*, *rbohD* and *rbohDF* with or without 24-hour treatment of 1 *μ* m CIF2. Brackets and arrowheads indicate Casparian strips and H_2_O_2_ production sites respectively. Scale bar = 500 nm. (E) Quantification of ROS production as number of dark pixels area (n = 23 – 44 sites from 5 roots of each condition). For quantification methods, see fig. S4.

In contrast to lignin polymerization, suberin might not require NADPH oxidase activity. It was shown previously that CIF2 triggers excess suberization in WT (Fig. S3B)^15^. We found excess suberin deposition in *rbohD* and *rbohF* upon peptide treatment, although a slight enhancement is already observed in *rbohF*. In *rbohDF*, excess suberin deposition is very strong, even without treatment, likely due to a strong activation of the surveillance system due to complete absence of a CS (Fig. S3B). These data indicate that CIF2-triggered suberin accumulation is not affected by NADPH oxidases and suggests existence of a ROS-independent branch of the SCHENGEN pathway that regulates this process (see below).

### SGN1 can directly activate RBOHF and RBOHD via phosphorylation

The above data strongly suggest a direct connection between the SGN3/SGN1 kinase module and the two NOX enzymes. We therefore conducted an *in vitro* kinase assay in order to ask whether SGN1 can directly phosphorylate the N-terminal cytoplasmic region of RBOHF and RBOHD. We found that recombinantly expressed TF-SGN1 could phosphorylate both the recombinant N-terminal part of RBOHF and RBOHD (Fig. 5A). Plant NOX regulation has been intensively studied and shown to be highly complex, requiring possibly interdependent activities of kinases, as well as small GTPases ^26^. We therefore tested whether SGN1 might be sufficient for activation of RBOHF activity in a cellular context. To do so, we made use of a heterologous reconstitution system in human HEK293T cells, which show very low endogenous ROS production and for which it had been previously demonstrated that plant NADPH oxidases can be expressed and their activation mechanism be studied ^27^. As a positive control the previously described calcium-dependent kinase complex of calcineurin B-like (CBL) interacting protein kinases 26 (CIPK26) and CBL1 was used and shown to be active (Fig. 5B). When expressing wild-type SGN1 in this cell line, we noticed that it did not activate RBOHF or RBOHD, but that it also did not localize to the plasma membrane as in plant cells (Fig. S5A). However, when we used the functional, constitutively plasma membrane-localized myrpalm-SGN1 version, a significant induction of ROS production was observed for RBOHF and to a lesser extent for RBOHD (Fig. 5B, S5B).

**Fig. 5.**
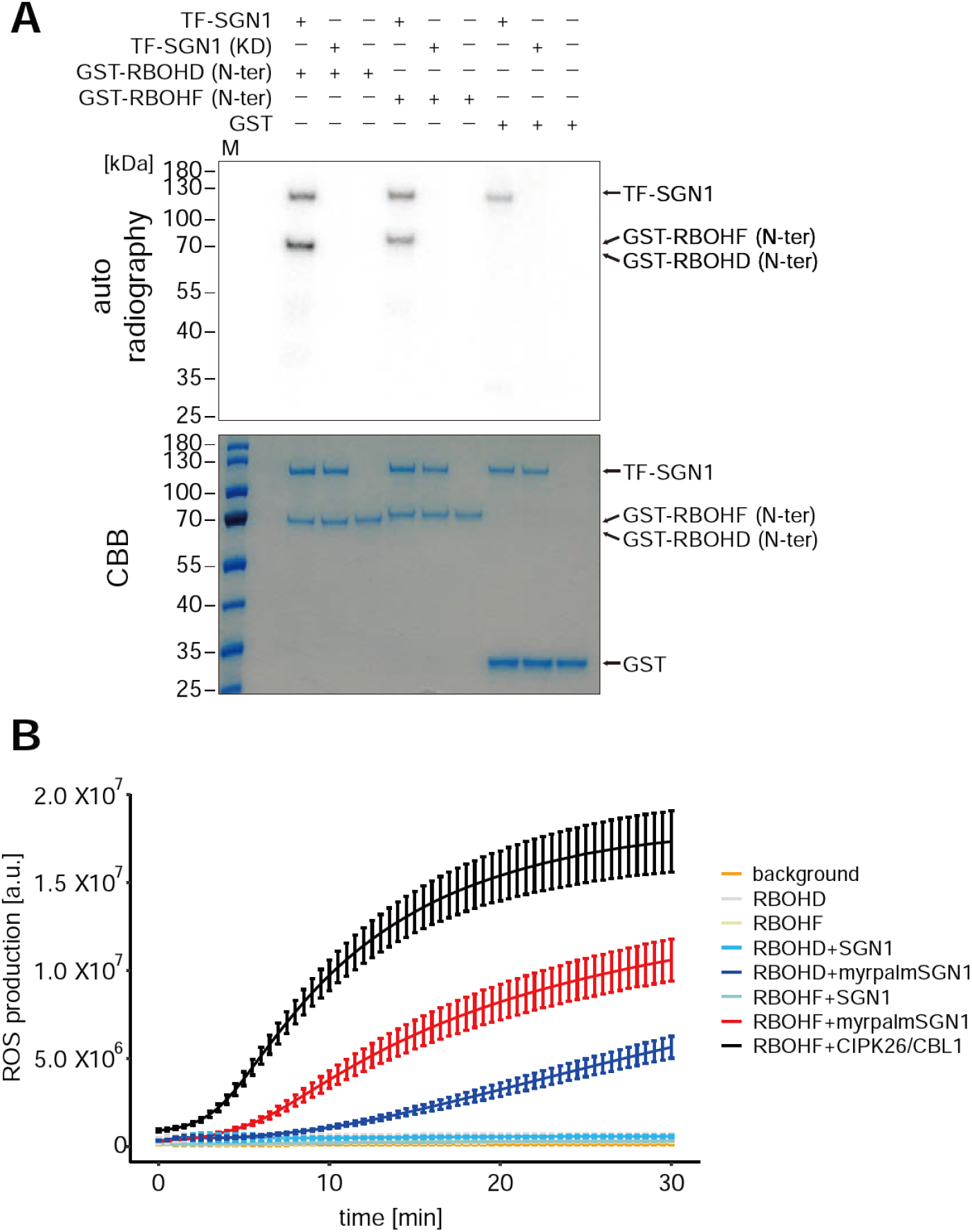
SGN1 directly activates NADPH oxidases in a cellular context. (A) *In vitro* kinase assay of TF-SGN1 against GST-N-terminal cytoplasmic domains of RBOHD or F. Experiments were done independently three times with similar results. (B) HEK293Tcell based NOX activation assay. Cells were transfected with the indicated plasmid combinations. The phosphatase inhibitor Calyculin A was added directly before the start of the measurements. Each data point represents the mean of six wells analyzed in parallel, bars indicate S.D.. Experiments were repeated three times (another set of results is shown in Fig. S5B).

### CIF peptide-induced CASP domain growth requires new protein synthesis, but not ROS production

The direct phospho-relay from SGN3 receptor, to SGN1 kinase to RBOHF and RBOHD outlined above draws a direct molecular connection from perception of a peptide hormone stimulus to cell wall lignification. Moreover, it accounts for both, the highly localized ROS production that we observe upon CIF stimulation, as well as the observation that localization of lignification is correlated with the site of active SGN3 signaling. Yet, the massive enhancement of lignification observed upon CIF stimulation (Fig3B), the increase of CASP accumulation and ectopic patch formation (Fig. S2A), as well as the non-localized formation of precocious and enhanced suberin (Fig. S3B,C), are additional outcomes of SGN3 pathway stimulation and should be driven by transcriptional changes. Indeed, some degree of transcriptional upregulation of CASP genes has been reported previously upon CIF1 treatment ^17^. CIF stimulation does not only lead to enhanced accumulation and ectopic patches of CASP1-GFP, absence of SGN3 signaling also leads to discontinuous CASP1-GFP signals that could be explained by insufficient amount of CASPs and other factors being produced during endodermal differentiation (Fig. 6A) ^15,18^. Neither the single, nor the double NADPH oxidase mutants display discontinuous CASP1-GFP signals, indicating that ROS production is not required for this aspect of the CIF/SGN3 pathway (Fig. 6A). In order to directly demonstrate that formation of a continuous CASP domain requires newly formed gene products, we treated our *cif1 cif2* double mutant with CIF2 peptide in presence or absence of protein synthesis inhibitor cycloheximide (CHX). CASP1-GFP signal strongly increased during 8 hours of CIF treatment, during which the discontinuous CASP1-GFP domains became continuous. This effect was abrogated by CHX treatment (Fig. 6 B,C, Movie S1).

**Fig. 6.**
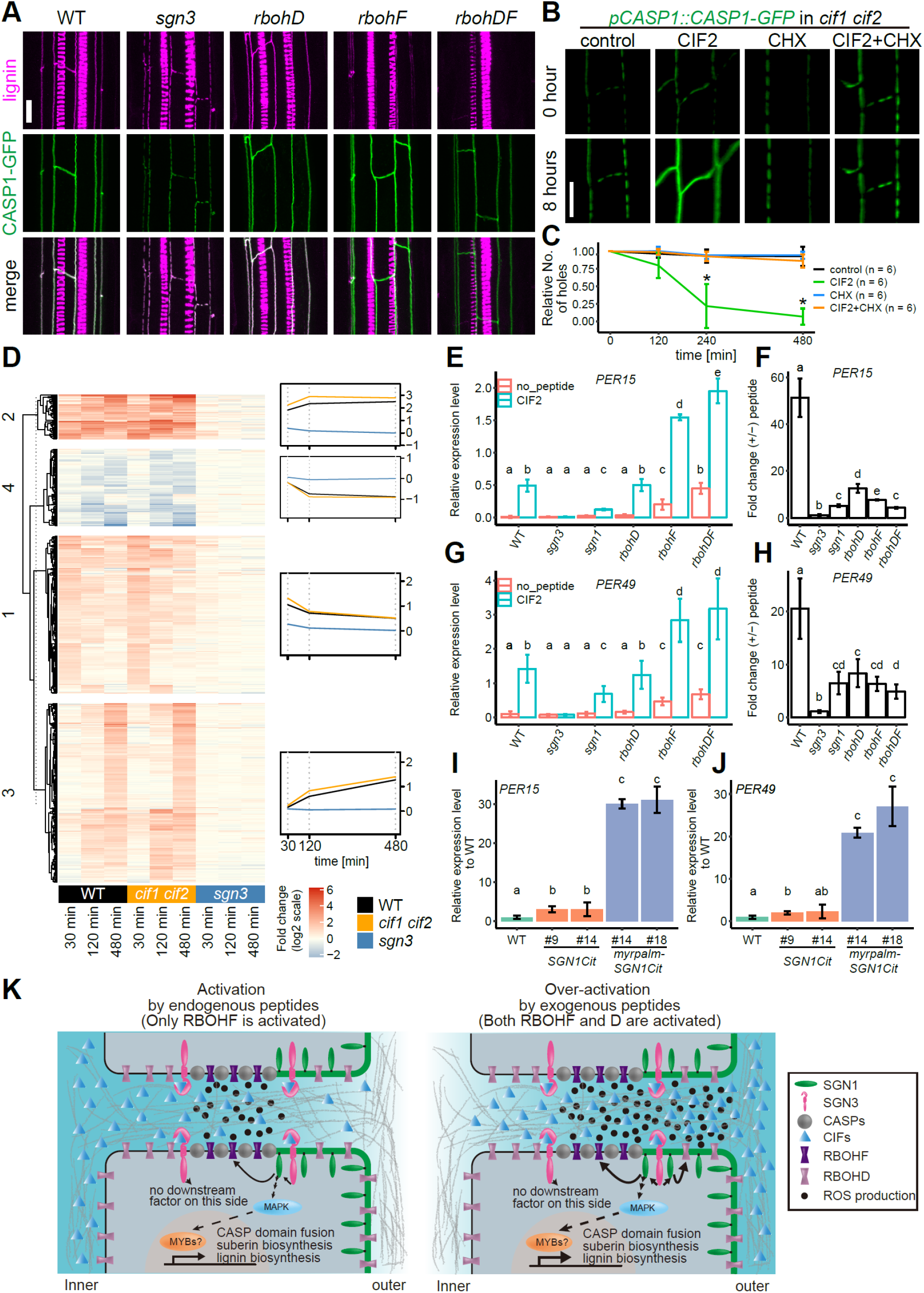
CIF2 induces large-scale transcriptional changes for cell wall remodeling. (A) CASP1-GFP and lignin deposition in WT, *sgn3*, *rbohD*, *rbohF* and *rbohDF*. CASP1-GFP and lignin (fuchsin) are presented in green and magenta respectively. Scale bar = 10 *μ* m. (B) Time lapse imaging of single- or cotreatment of CIF2 with cycloheximide (CHX) on CASP1-GFP in *cif1 cif2.* Scale bar = 10 *μ* m (See also supplemental movie 1). (C) Quantification of (B). Relative numbers of holes in CASP1-GFP domain after single- or co-treatment with CIF2 or CHX from the pictures in (B). Bars are S.D. * indicates statistical significance from all other conditions (p<0.01) after one-way ANOVA and Tukey test. (D) Fold-change of 930 genes (p<0.05 and log2(Fold change) ≥1 or ≤ −1 at least one time point in one genotype) after CIF2 treatment at indicated time points in WT, *cif1,2* and *sgn3*. Degree of the fold changes are shown in color code as indicated. (E) Relative expression levels of *PER15* to *CLATHRIN* control in each genotype with or without 2-hour CIF2 treatment. Bars are S.D. (n=3). Different characters indicate statistically significant differences (p < 0.01, ANOVA and Tukey test). (F) Fold changes of *PER15* in each genotype with or without 2-hour CIF2 treatment. Bars are S.D. (n=3). Different characters indicate statistical significance differences (p < 0.01, ANOVA and Tukey test). (G) Relative expression levels of *PER49* to *CLATHRIN* control in each genotype with or without 2-hour CIF2 treatment. Bars are S.D. (n=3). Different characters indicate statistically significant differences (p < 0.01, ANOVA and Tukey test). (H) Fold changes of *PER49* in each genotype with or without 2-hour CIF2 treatment. Bars are S.D. (n=3). Different characters indicate statistically significant differences (p < 0.01, ANOVA and Tukey test). (I) Relative fold changes of *PER15* in *pCASP1∷SGN1-Cit* and *pCASP1∷myrpalm-SGN1-Cit* compared to the expression level in WT. Bars are S.D. (n=3). Different characters indicate statistically significant differences (p < 0.01, ANOVA and Tukey test). (J) Relative fold changes of *PER49* in *pCASP1∷SGN1-Cit* and *pCASP1∷myrpalm-SGN1-Cit* compared to the expression level in WT. Bars are S.D. (n=3). Different characters indicate statistically significant differences (p < 0.01, ANOVA and Tukey test). (K) Schematic overview of how localized ROS production and gene activation are integrated in the SCHENGEN pathway and how localized ROS production is achieved upon endogenous stimulation (left) and by exogenous application with high amounts of ligand (right). “inner” designates the stele-facing endodermal surface, “outer”, the cortex-facing surface.

### The NADPH oxidase-independent branch of the CIF/SGN3 pathway is associated with MAP kinase stimulation and causes strong activation of gene expression

In pattern-triggered immune receptor signaling, gene activation is thought to depend in large parts on activation of Mitogen-activated protein (MAP) kinases ^28^. We therefore tested whether CIF-treatment leads to MAP kinase phosphorylation and indeed found that CIFs can induce MAP kinase phosphorylation in a SGN3-dependent manner, further extending the molecular parallels between immune receptor signaling and the CIF/SGN3 pathway and suggesting that gene induction in the CIF pathway might equally depend on MAP kinase signaling (Fig. S6B).

We then undertook an RNA profiling of seedling roots at different time points (30, 120 and 480 min) after CIF stimulation, using wild-type, *cif1 cif2* and *sgn3* mutants as genotypes. 930 genes were found to be differentially expressed across any of the genotypes and time points combined, using a stringent cut-off (adj.-pval. <= 0.05; logFC >= 1 or logFC<=−1) (Table S1). After normalization of batch effects, the three replicates clustered closely with a large degree of variance explained by the peptide treatment in wild-type and *cif1 cif2* (Fig. S6A). Wild-type and *cif1 cif2* samples showed nearly identical responses (co-relation efficiency values were 0.90, 0.89, 0.94 at 30, 120, 480 min respectively), with *cif1 cif2* displaying a slightly stronger overall amplitude (Fig. 6D). Importantly, *sgn3* had virtually no differentially expressed genes across treatments, indicating the specificity of the CIF responses and further corroborating that SGN3 is the single, relevant receptor for CIF responses in roots (Fig. 6D, S6A, Table S1). Our data extends on the previous data of Nakayama et al., by showing that all 5 *CASP* genes are differentially regulated upon CIF-treatment (Fig. S6C). Moreover, we observed upregulation of *MYB36*, a central transcription factor for Casparian strip formation and CASP expression, thus potentially accounting for the increases in *CASP1-4* expression ^20,29^(Fig. S6C). The fact that CIF stimulates *MYB36* expression also is consistent with the recent report that CIF treatment can enhance ectopic endodermal differentiation, driven by overexpression of the SHR transcription factor ^30^.

Of the four response clusters defined by k-mean clustering, the two largest clusters (1 and 3) group early response genes (cluster 1) and later response genes (cluster 3). Cluster 2 groups a smaller fraction of genes showing strong and largely sustained responses over the time scale of the experiment, while cluster 4 groups a minor fraction of genes downregulated at later time points. GO term analysis indicates that many of the most significant, overrepresented terms in the “early response” (1) and “strong and sustained” (2) gene clusters are related to immune and defense responses (response to chitin, bacterium, callose, etc.), as well as responses to oxidative stress (Table S2). The top GO categories of the late response cluster (3) were suberin/cutin biosynthesis, supporting our observation that suberin accumulation is a late effect of CIF stimulation (Fig. S6D, S3B,C)^15^. Additional, enriched terms were related to oxidative stress and cell wall remodeling, nicely fitting the CIF2-induced ROS production we observed. Intriguingly, the cluster showing late downregulation of gene expression contains many overrepresented GO terms related to a wide variety of transport processes, reaching from water, nitrate and ammonium transport, to primary and secondary metabolite or hormone transport (Fig. 6D, Table S2). We interpret this as a consequence of CIF-induced lignification and suberization, which must profoundly impact endodermal ability for shuttling organic and inorganic compounds, as well as signaling molecules between stele and cortical root cell layers ^10^.

While GO terms related to lignification were also overrepresented, they were not as highly ranked as expected from the strong induction of lignification upon CIF treatment. A number of reasons might account for this: Firstly, GO terms for pathogen, defense and oxidative stress responses, are all bound to overlap with and contain genes that mediate lignification. Moreover, already elevated levels of developmental, lignin-related gene expression in the endodermis might lead to less-pronounced fold-changes. Finally, the multiple roles for lignin biosynthetic genes in secondary metabolism further obscure clear categorization. Nevertheless, upregulation of lignin biosynthesis by CIF-treatment became evident when clustering a set of laccases and peroxidases, the two main families of enzymes implicated in lignin polymerization in the apoplast ^7^. A significant number of both are strongly upregulated (Fig. S6E,F). Moreover, *MYB15*, a transcription factor shown to be involved in stress and MAMP(microbe-associated molecular pattern)-induced lignification ^31^ is about 6-fold induced after 30 min and about 17-fold after 120 min in both treated wild-type and *cif1 cif2*, nicely correlating with the upregulation of peroxidases and laccases (Fig. S6E,F, Table S1). We then made use of two of the most highly differentially expressed genes upon CIF2-treatment (*PER15* and *PER49*) in order to establish whether SGN1 and the NADPH oxidases are also required for gene regulation downstream of SGN3 activation. In qPCR analysis both peroxidase genes still showed a slight, but significant upregulation upon CIF-treatment in the *sgn1* mutant (compared to the complete absence of response in *sgn3*) (Fig. 6E-H), clearly indicating that, as for ROS production, SGN1 is an important, yet not absolutely required, downstream component in CIF-regulated gene expression. When directly plotting peroxidase fold-inductions in *rbohF*, *rbohD* and double mutant, a seemingly strong attenuation was observed (Fig. 6F,H). However, when independently plotting gene expression in treated and untreated conditions (Fig. 6E,G), this lower fold-induction is largely explained by an already significant basal increase in marker gene expression in *rbohF* and *rbohD rbohF* double mutants. Indeed, both mutants display an absence of Casparian strips, which should lead to an endogenous stimulation of the SGN3 pathway, nicely supporting the barrier surveillance model. Absolute expression of both marker genes in the NADPH oxidase mutants is in effect higher than in wild-type after CIF treatment, leading us to conclude that ROS production and gene activation in response to CIF are two independent branches of this signaling pathway, with the branching occurring downstream of the SGN1 kinase. We finally wanted to know whether the mis-localization of the SGN1 kinase reported initially is only affecting CIF/SGN3 signaling at the plasma membrane, or whether it also affects gene activation. We found that both *PER15* (Fig. 6I) and *PER49* (Fig. 6J) are dramatically upregulated in the non-polar SGN1 lines, independent of exogenous peptide application, strongly corroborating our model whereby the SCHENGEN pathway function crucially depends on the correct subcellular localization of its downstream kinase.

## DISCUSSION

The data presented here delineate an entire signaling pathway. Previously, SGN3 had been established as the receptor for CIF1 and 2, but its connection to SGN1 had exclusively been based on genetic evidence. Here we show that SGN3, SGN1 and RBOHD/F are, biochemically and functionally, part of a signal transduction chain, leading directly from localized peptide perception to localized ROS production and lignification (Fig. 6K). Moreover, the pathway branches downstream of SGN1, leading to MAP kinase activation and stimulation of gene expression. Some of the most strongly induced genes being peroxidases and laccases that would further enhance and sustain lignification. The SCHENGEN pathway therefore elegantly integrates fast, plasma membrane-based responses that maintain positional information (ROS and lignin is produced close to where the ligand is perceived) and slower, gene expression-based responses, which have lost positional information, but would allow to enhance and maintain the ROS-burst-controlled activation of peroxidases (Fig. 6K).

The SCHENGEN pathway bears a striking overall resemblance to well-established signaling pathways for perception of MAMPs. In MAMP perception, structurally similar receptor kinases, such as FLS2 or EFR bind to microbial patterns and transduce this signal through kinases of the RLCKVII family, such as BIK1 or PBLs, which are homologs of SGN1 ^32^. BIK1 in turn was shown to phosphorylate RBOHD, driving the well-described MAMP-induced ROS burst^33,34^. Moreover, it has recently been shown that kinases of the RLCKVII family directly phosphorylate MAPKKKs, which now mechanistically explains how MAMP perception induces MAP kinase phosphorylation ^35^. It is thus tempting to speculate that the SCHENGEN pathway represents an ancient neo-functionalization of an immune receptor pathway. Indeed, this has been proposed recently and it was pointed out that the closest receptor homologs to SGN3 and GSO2 are PEPR1 and PEPR2 ^36^. The latter are receptors to an endogenous plant peptides (AtPEPs), whose activities resemble that of MAMPs and are best thought of as “phytocytokines”, i.e. agents able to induce an immune-like response in cells that have not yet encountered MAMPs ^37^. Such an original phytocytokine might then have been neo-functionalized to induce pre-formed defensive cell wall barriers in a developmental context, in the absence of any actual biotic or abiotic stress. MAMP perception and MAMP-induced ROS production was found already in mosses and clearly precedes the wide-spread adoption of lignin as a major, cell wall reinforcing polymer ^38,39^. Moreover, lignin was speculated to originate from phenylpropanoid-derived defense compounds, making it plausible that immune signaling pathways have been at the origin of developmental regulation of lignification. The intriguing innovation of the SCHENGEN pathway would then reside in the subcellular arrangement of its signaling components, especially that of SGN1. Indeed, we demonstrate that polar localization of SGN1 to the outside of the Casparian strip domain is crucial for barrier function - since its simple mis-localization to the inside leads to ligand-and receptor-dependent overlignification at inner cell corners. Thus, the only feature that arrests signaling in wild-type is the formation of a lignified diffusion barrier in the cell wall, preventing access of the stele-produced peptides to SGN3 receptors at the outer domain - as the only population able to stimulate the polar SGN1 kinase. Since we have shown here that a main read-out of the SCHENGEN pathway is cell wall lignification itself, this designs a fascinating, spatial negative-feedback loop, in which SCHENGEN pathway-stimulated lignification feedback-regulates itself once enough lignin has been produced to form a tight diffusion barrier and to fully prevent further CIF peptide penetration to the outside.

It will be important to further describe the extent to which the SCHENGEN pathway uses components of MAMP signaling (as for example the requirement for SERK-family co-receptors, which has not been demonstrated yet for SGN3), but we propose that our findings on the SCHENGEN pathway function can be of considerable interest for MAMP receptor signaling. The particular, restricted spatial overlap of receptor and downstream kinase has allowed us to visualize that, even if NADPH oxidases are non-localized (as in the case of RBOHD), ROS can be locally produced in a micrometer-scale region at the plasma membrane, through localized receptor stimulation. In MAMP receptor signaling, the non-localized nature of both MAMP receptor and NADPH oxidase would not have allowed to visualize this unexpected degree of spatial control. Yet, during actual microbial infections, highly-localized receptor stimulation and ROS production might indeed occur and be relevant for the outcome of the immune response. In addition, while it is established that MAMP-stimulated ROS production is an important part of the immune response, its direct molecular downstream action is not well understood. Lignin production has long been associated with immune responses, as well as responses to cell wall damage, yet a direct molecular connection from MAMP perception to lignification has rarely been drawn ^31^. It will be intriguing to investigate whether and how much MAMP-induced ROS production is actually used for lignification during defense and to which degree this explains the importance of ROS in plant defense responses.

The SCHENGEN pathway might not be limited to regulation of lignified diffusion barriers, as it has been shown that it is also important in the formation of the embryonic cuticle. The peptide ligand used in this context has not been identified and it remains unclear whether an equally precise barrier surveillance mechanism is also acting to ensure separation of endosperm and embryo during embryonic cuticle formation ^36^. In this context, the SCHENGEN pathway might be used to drive production and deposition of cutin instead of lignin and suberin as in the case of the endodermis. In the future, it will be fascinating to investigate whether the SCHENGEN pathway is of even broader developmental significance and to understand the molecular basis that enables its distinct, organ and cell-type specific activities.

## Acknowledgments

We thank the Central Imaging Facility (CIF), Genome Technology Facility (GTF), particularly Sandra Calderon, and Electron Microscopy facility (EMF) of the University of Lausanne for expert technical support. We also thank Hiroko Uchida for expert graphical support.

## Author contributions

S.F. and N.G. conceived the project. S.F., T.G.A and N.G. designed the experiments. S.F., D.D.B., K.H.E, P.K, T.G.A, V.D.T, A.P., P.M., R.U, V.G.D. and A.C. performed the experimental work. D.D.B., T.G.A., E.S.S. and J.D. performed image quantification and RNA-seq analysis. S.F., and N.G. wrote the manuscript. G.I., J.K. and all other authors revised the manuscript and were involved in the discussion of the work.

## Data and materials availability

All data to support the conclusions of this manuscript are included in the main text and supplementary materials.

## Supplementary Materials

Materials and Methods

Figures S1-S6

Tables S1-S2

Movies S1

References (*40-61*)

## References

1. Kärkönen, A. & Kuchitsu, K. Reactive oxygen species in cell wall metabolism and development in plants. Phytochemistry 112, 22–32 (2015).

2. Zipfel, C. Plant pattern-recognition receptors. Trends in Immunology 35, 345–351 (2014).

3. Qi, J., Wang, J., Gong, Z. & Zhou, J.-M. Apoplastic ROS signaling in plant immunity. Current Opinion in Plant Biology 38, 92–100 (2017).

4. Boisson-Dernier, A. et al. ANXUR Receptor-Like Kinases Coordinate Cell Wall Integrity with Growth at the Pollen Tube Tip Via NADPH Oxidases. PLoS Biology 11, e1001719 (2013).

5. Monshausen, G. B., Bibikova, T. N., Messerli, M. A., Shi, C. & Gilroy, S. Oscillations in extracellular pH and reactive oxygen species modulate tip growth of Arabidopsis root hairs. Proceedings of the National Academy of Sciences 104, 20996–21001 (2007).

6. Barbosa, I. C. R., Rojas-Murcia, N. & Geldner, N. The Casparian strip—one ring to bring cell biology to lignification? Current Opinion in Biotechnology 56, 121–129 (2019).

7. Liu, C.-J. Deciphering the Enigma of Lignification: Precursor Transport, Oxidation, and the Topochemistry of Lignin Assembly. Molecular Plant 5, 304–317 (2012).

8. Lee, Y., Rubio, M. C., Alassimone, J. & Geldner, N. A Mechanism for Localized Lignin Deposition in the Endodermis. Cell 153, 402–412 (2013).

9. Geldner, N. The endodermis. Annu Rev Plant Biol 64, 531–558 (2013).

10. Barberon, M. & Geldner, N. Radial transport of nutrients: the plant root as a polarized epithelium. Plant Physiol. 166, 528–537 (2014).

11. Alassimone, J., Naseer, S. & Geldner, N. A developmental framework for endodermal differentiation and polarity. Proc. Natl. Acad. Sci. U.S.A. 107, 5214–5219 (2010).

12. Naseer, S. et al. Casparian strip diffusion barrier in Arabidopsis is made of a lignin polymer without suberin. Proc. Natl. Acad. Sci. U.S.A. 109, 10101–10106 (2012).

13. Hosmani, P. S. et al. Dirigent domain-containing protein is part of the machinery required for formation of the lignin-based Casparian strip in the root. Proc. Natl. Acad. Sci. U.S.A. 110, 14498–14503 (2013).

14. Roppolo, D. et al. A novel protein family mediates Casparian strip formation in the endodermis. Nature 473, 380–383 (2011).

15. Doblas, V. G. et al. Root diffusion barrier control by a vasculature-derived peptide binding to the SGN3 receptor. Science 355, 280–284 (2017).

16. Alassimone, J. et al. Polarly localized kinase SGN1 is required for Casparian strip integrity and positioning. Nature Plants 2, 16113 (2016).

17. Nakayama, T. et al. A peptide hormone required for Casparian strip diffusion barrier formation in Arabidopsis roots. Science 355, 284–286 (2017).

18. Pfister, A. et al. A receptor-like kinase mutant with absent endodermal diffusion barrier displays selective nutrient homeostasis defects. eLife 3, e03115 (2014).

19. Kalmbach, L. et al. Transient cell-specific EXO70A1 activity in the CASP domain and Casparian strip localization. Nature Plants 3, 17058 (2017).

20. Kamiya, T. et al. The MYB36 transcription factor orchestrates Casparian strip formation. Proc. Natl. Acad. Sci. U.S.A. 112, 10533–10538 (2015).

21. Li, B. et al. Role of LOTR1 in Nutrient Transport through Organization of Spatial Distribution of Root Endodermal Barriers. Current Biology 27, 758–765 (2017).

22. Vermeer, J. E. M., Munster, E. B. V., Vischer, N. O. & Gadella, T. W. J. Probing plasma membrane microdomains in cowpea protoplasts using lipidated GFP-fusion proteins and multimode FRET microscopy. Journal of Microscopy 214, 190–200 (2004).

23. Kim, T.-W., Guan, S., Burlingame, A. L. & Wang, Z.-Y. The CDG1 Kinase Mediates Brassinosteroid Signal Transduction from BRI1 Receptor Kinase to BSU1 Phosphatase and GSK3-like Kinase BIN2. Molecular Cell 43, 561–571 (2011).

24. Lu, D. et al. A receptor-like cytoplasmic kinase, BIK1, associates with a flagellin receptor complex to initiate plant innate immunity. Proc. Natl. Acad. Sci. U.S.A. 107, 496–501 (2010).

25. Bestwick, C. S., Brown, I. R., Bennett, M. H. & Mansfield, J. W. Localization of hydrogen peroxide accumulation during the hypersensitive reaction of lettuce cells to Pseudomonas syringae pv phaseolicola. The Plant Cell 9, 209–221 (1997).

26. Kadota, Y., Shirasu, K. & Zipfel, C. Regulation of the NADPH Oxidase RBOHD During Plant Immunity. Plant and Cell Physiology 56, 1472–1480 (2015).

27. Han, J.-P. et al. Fine-tuning of RBOHF activity is achieved by differential phosphorylation and Ca2+ binding. New Phytologist 221, 1935–1949 (2019).

28. Dodds, P. N. & Rathjen, J. P. Plant immunity: towards an integrated view of plant–pathogen interactions. Nature Reviews Genetics 11, 539–548 (2010).

29. Liberman, L. M., Sparks, E. E., Moreno-Risueno, M. A., Petricka, J. J. & Benfey, P. N. MYB36 regulates the transition from proliferation to differentiation in the *Arabidopsis* root. Proceedings of the National Academy of Sciences 112, 12099–12104 (2015).

30. Drapek, C. et al. Minimum requirements for changing and maintaining endodermis cell identity in the Arabidopsis root. Nature Plants 4, 586–595 (2018).

31. Chezem, W. R., Memon, A., Li, F.-S., Weng, J.-K. & Clay, N. K. SG2-Type R2R3-MYB Transcription Factor MYB15 Controls Defense-Induced Lignification and Basal Immunity in Arabidopsis. The Plant Cell 29, 1907–1926 (2017).

32. Liang, X. & Zhou, J.-M. Receptor-Like Cytoplasmic Kinases: Central Players in Plant Receptor Kinase–Mediated Signaling. Annual Review of Plant Biology 69, 267–299 (2018).

33. Kadota, Y. et al. Direct Regulation of the NADPH Oxidase RBOHD by the PRR-Associated Kinase BIK1 during Plant Immunity. Molecular Cell 54, 43–55 (2014).

34. Li, L. et al. The FLS2-Associated Kinase BIK1 Directly Phosphorylates the NADPH Oxidase RbohD to Control Plant Immunity. Cell Host & Microbe 15, 329–338 (2014).

35. Bi, G. et al. Receptor-Like Cytoplasmic Kinases Directly Link Diverse Pattern Recognition Receptors to the Activation of Mitogen-Activated Protein Kinase Cascades in Arabidopsis. The Plant Cell 30, 1543–1561 (2018).

36. Creff, A. et al. A stress-response-related inter-compartmental signalling pathway regulates embryonic cuticle integrity in Arabidopsis. PLOS Genetics 15, e1007847 (2019).

37. Gust, A. A., Pruitt, R. & Nürnberger, T. Sensing Danger: Key to Activating Plant Immunity. Trends in Plant Science 22, 779–791 (2017).

38. Bressendorff, S. et al. An Innate Immunity Pathway in the Moss *Physcomitrella patens*. The Plant Cell 28, 1328–1342 (2016).

39. Weng, J.-K. & Chapple, C. The origin and evolution of lignin biosynthesis: Tansley review. New Phytologist 187, 273–285 (2010).

