## Supplemental information for "ROS production by localized SCHENGEN receptor module drives lignification at subcellular precision"

### Materials and Methods

**Plant material and growth conditions**. For all experiments, *Arabidopsis thaliana* (ecotype Columbia) was used. The T-DNA tagged lines, *sgn1-2* (SALK_055095C) and *sgn3-3* (SALK_043282) were obtained from NASC(*16*, *18*). *rbohD dSpm*, *rbohF dSpm* and *rbohD dSpm rbohF dSpm* are described in (*40*, *41*). *esb1-1* was kindly shared by Prof. David Salt's group (*42*, *43*). The *cif1-2 cif2-2* double mutant was generated by CRISPR-Cas9 method (*44*) using 5'- gctttggtttaggactggag -3' as a protospacer sequence to target both CIF1 and CIF2 loci.

*pCASP1::CASP1–GFP* marker line was crossed into *sgn1-2*, *sgn3-3* or *cif1-2 cif2-2*. The *pCASP1::myrpalm SGN1* construct was independently transformed into each mutant background using the floral dip method(*45*). For observations and histological analysis, seeds were kept for 2 days at 4°C in the dark for stratification, then grown for 5 days at 22°C under 16 h light/8 h dark vertically on solid half-strength Murashige–Skoog (MS) medium. Cycloheximide (Sigma) and Diphenyleneiodonium chloride (Sigma) were prepared as 50 mM water stock solution and as 20 mM DMSO stock solution.

#### Plasmid construction

To generate myrpalm SGN1, GGCFSKK (5'- GGAGGATGCTTCTCTAAGAAG -3') (*22*) was added to SGN1 cDNA right after the start codon by inverse PCR. This SGN1 variant was combined with pCASP1 and mCitrine coding sequence by LR clonase (Thermo Fisher Scientific). Coding sequences of RBOHF 2-383 a.a. and RBOHD 1-376 a.a. were inserted between BamHI and XhoI sites of a modified pET24(+) vector, in which GST (glutathione S-transferase) was added on N-terminus. For constructing a GST-SGN3 kinase domain vector, a cDNA fragment coding for 899 a.a. to 1249 a.a. of SGN3 was fused into pDEST15 by Gateway LR reaction. TF-SGN1 was as described previously (*16*). Kinase-dead mutations in SGN1 (K134E) and SGN3 (K979E) were introduced by site-directed mutagenesis.

#### Protein expression in bacteria and purification

GST or Trigger factor (TF) proteins were expressed from pGex6p-1 or pCold-TF vectors in BL21 (DE3) CodonPlus RIL and purified in a glutathione sepharose column (Thermo Fischer Scientific) or a Co-Sepharose column (Clontech) according to the manufacturer's protocols. GST-RBOHD N-ter and GST-RBOHF N-ter were produced and purified as described (*46*). Bacterially expressed GST-SGN3 kinase domain (WT or kinase-dead) in BL21 (DE3) CodonPlus RIL was purified with a glutathione sepharose column following the manufacture's protocol. For TF-SGN1 (WT or kinase-dead), we followed the method in the previous report (*16*). The buffer of all proteins was changed to 50 mM HEPES-KOH pH7.5 and 1 mM DTT with PD-10 or NAP-5. Purified kinase protein solutions were mixed with glycerol to 30% [v/v] as final concentration and kept at -20°C until use.

#### *In vitro* kinase assay

*In vitro* kinase assay was done as described previously (*16*) with small modifications. Purified 250 ng GST-SGN3 kinase domain was incubated with 250 ng TF-SGN1 or TF proteins in reaction buffer (50 mM HEPES-KOH pH7.5, 1 mM MnCl2. 1 mM DTT, 1 mM ATP, 185,000 Bq [γ-$^{32}$P] ATP) at 30°C for 30 min. 250 ng TF-SGN1 and 1 μg GST or 500 ng GST-N-terminal cytoplasmic domains of RBOHD or F were treated in the same way as above. The reaction was stopped by adding 4 x LDS sample buffer (Invitrogen) and heating at 75°C for 10 min. The samples were separated on 4-12 % gradient Nu-PAGE gels or 10 % Nu-PAGE gels (Invtirogen). After drying the gels, signal was detected using Typhoon FLA7000 (GE Healthcare).

#### Western blotting

Plants were grown in liquid MS medium supplemented with 0.5% sucrose with cell strainer for 5 days under long day (18 h light /6 h dark) condition after 2-day vernalization at 4°C. Hydroponically grown seedlings were transferred into fresh liquid MS medium containing 0.5% sucrose with or without 1 μM CIF2 peptide and incubated for 15 min. The seedlings were immediately frozen in liquid nitrogen and grinded by TissueLyser II (Qiagen). Extraction buffer (20 mM Na phosphate pH 7.4, , 150 mM NaCl, 1 mM EDTA,, 0.1% Tween, 50 mM β glycerophosphate, 0.5 mM PMSF, 100 μM sodium orthovanadate, 5 mM Na fluoride, Complete cocktail) was added to the frozen samples. The samples were briefly mixed by vortex and centrifuged at 15,000 rpm for 15 min at 4°C. The supernatant was transferred in new tubes and measured the concentration by Bradford method (Thermo Fischer Scientific). Equal amount of protein was loaded (20 μg / lane) and separated in 10% acrylamide gel (Eurogentec). After the electrophoresis, the separated proteins were transferred onto nitrocellulose membrane (Amersham, GE Healthcare) by XCell SureLock (Invitrogen) and stained with Ponceau S to show the loading control. The blot was incubated in blocking buffer (3% skim milk in TBS) for 1 hour and probed with anti-phospho-p42, p44 antibody (1:1000 dilution, Cell signaling technologies #9101) in 0.5% skim milk in TBS-0.1% Tween for overnight at 4°C. After washing the blot three times with TBS-0.1% Tween, the blot was treated with anti-rabbit secondary antibody (1:30,000 dilution, Agrisera) in 0.5% skim milk in TBS-0.1% Tween for 1 hour and washed three times with TBS-0.1% Tween. Signals were detected on X-ray film with SuperSignal West Femto Kit (Thermoscientific). For the MPK6 detection, the membrane was stripped (Pierce), incubated in blocking buffer for 1 hour and probed with anti-MPK6 antibody (1:10000 dilution, Sigma) in blocking buffer for 1 hour at room temperature. After washing the blot three times with TBS-0.1% Tween, the membrane was treated with anti-rabbit secondary antibody (1:30000, Agrisera) in blocking buffer and washed again three times. Signal were detected with SuperSignal West Femto kit (Thermo Scientific).

**Confocal microscopes**

Confocal pictures were obtained using Leica SP8 or Zeiss LSM 880 confocal microscopes. The excitation and detection window settings to obtain signal as follows: When using Leica SP8 (excitation, detection window), GFP (488 nm, 500–550 nm), mCitrine, (514 nm, 518–560 nm), mCitrine/mCherry, (514 nm and 594 nm, 518–560 nm, 600–650 nm, sequential scan), when using Zeiss LSM880 (excitation, detection window), Calcofluor White (405 nm, 425-475 nm), Basic Fuchsin (561 nm, 570-650 nm) and Fluorol Yellow (488 nm, 500-550 nm).

**PI assay**

PI assay was done as described previously (*8*) with small changes. Seedlings were incubated in water containing 10 $\mu$g /mL PI for 10 min and transferred into fresh water. The number of endodermal cells were scored using a Zeiss LSM 700 confocal microscope (excitation 488 nm, SP640, split at 570 nm) from onset of cell elongation (defined as endodermal cell length being more than two times than width in the median, longitudinal section) until PI could not penetrate into the stele.

**Scoring discontinuities in CS domain**

CASP1-GFP signal was obtained by Leica SP8 as 1 µm step z-stack images from 5-day old seedlings. After the images were projected as maximum projections, total lengths of CS domains were measured in maximum projections and the number of discontinuities in the domain was counted manually.

**Lignin and cell wall staining**

ClearSee staining coupled cell wall staining were performed as described in recent publications (*47*, *48*). Briefly, 7-8 five-day-old seedlings were fixed in 3 mL 1 x PBS containing 4% para-formaldehyde for 1 hour at room temperature in 12-well plates and washed twice with 3 mL 1 x PBS. Following fixation, the seedlings were cleared in 3 mL ClearSee solution under gentle shaking. After overnight clearing, the solution was exchanged to new ClearSee solution containing 0.2% Fuchsin and 0.1% Calcofluor White for lignin and cell wall staining respectively. The dye solution was removed after overnight staining and rinsed once with fresh ClearSee solution. The samples were washed in new ClearSee solution for 30 min with gentle shaking and washed again in another fresh ClearSee solution for at least one overnight before observation.

**Methanol-based Fluorol Yellow staining of Arabidopsis root suberin**

Vertically grown 5-day old seedlings were incubated in methanol for three days at room temperature. The cleared seedlings were transferred to a freshly prepared solution of Fluorol Yellow 088 (0.01%, in methanol) and incubated for 1 hour. The stained seedlings were rinsed shortly in methanol and transferred to a freshly prepared solution of aniline blue (0.5%, in methanol) for counterstaining. Finally, the seedlings were washed for 2-3min in water and transferred to a chambered coverglass (Thermo Scientific), covered with a piece of 1% half strength MS agar and imaged using a Zeiss LSM 880 confocal microscope as described above.

#### Detection of $H_2O_2$ production sites in situ using transmission electron microscopy

Visualization of $H_2O_2$ production sites around Casparian strip was done by cerium chloride method as described previously (*8*, *49*) with some modifications. 4-day-old Arabidopsis seedlings were transferred onto fresh 1/2 MS solid medium with or without 1 $\mu$M CIF2 and incubated for 24 hours. After the peptide treatment, the seedlings were incubated in 50 mM MOPS pH7.2 containing 10 mM $CeCl_3$ for 30 min. After incubation with $CeCl_3$, seedlings were washed twice in MOPS buffer for 5 min and fixed in glutaraldehyde solution (EMS, Hatfield, PA) 2.5% in 100 mM phosphate buffer (pH 7.4) for 1 hour at room temperature. Then, they were post-fixed in osmium tetroxide 1% (EMS) with 1.5% of potassium ferrocyanide (Sigma, St. Louis, MO) in phosphate buffer for 1 hour at room temperature. Following that, the plants were rinsed twice in distilled water and dehydrated in ethanol solution (Sigma) at gradient concentrations (30% 40 min; 50% 40 min; 70% 40 min; two times (100% 1 hour). This was followed by infiltration in Spurr resin (EMS) at gradient concentrations (Spurr 33% in ethanol, 4 hours; Spurr 66% in ethanol, 4 hours; Spurr two times (100% 8 hours) and finally polymerized for 48 hours at 60°C in an oven. Ultrathin sections 50 nm thick were cut transversally at 1.3 ± 0.1 mm from the root tip, on a Leica Ultracut (Leica Mikrosysteme GmbH, Vienna, Austria) and picked up on a copper slot grid 2x1 mm (EMS) coated with a polystyrene film (Sigma). Micrographs were taken with a transmission electron microscope FEI CM100 (FEI, Eindhoven, The Netherlands) at an acceleration voltage of 80kV and 11000x magnification (pixel size of 1.851nm, panoramic of 17x17 pictures), exposure time of 800ms, with a TVIPS TemCamF416 digital camera (TVIPS GmbH, Gauting, Germany) using the software EM-MENU 4.0 (TVIPS GmbH, Gauting, Germany). All the picture were taken using the same beam intensity, and panoramic aligned with the software IMOD (*50*). For quantification, thresholding of cerium precipitate on pictures were scored using IMOD software (*50*). Briefly, all pictures subjected to the quantification was normalized to one picture, a section from non-treated wild type seedlings, by comparing the grey value of plastids in pericycle cells. After normalization, cell wall spaces from the beginning of Casparian strip in pericycle side to a cortex side between endodermal cells were selected and one identical threshold setting was applied to all pictures in order to highlight signals around Casparian strips. The values were shown as a percentage of thresholded pixels to selected area.

#### ROS production assay in HEK cells

Measurements of RBOHD and RBOHF activity were performed as described in (*27*) with some modifications. RBOHD or RBOHF were transiently expressed in HEK293T cells with or without coexpression of SGN1 or myrpalm SGN1. HEK293T cells were cultivated at 5 % $CO_2$ and 37 ° C in Dulbecco's modified Eagle's medium (DMEM) mixture F-12HAM (Sigma) enriched with 10% fetal bovine serum. Before transfection, lysine coated white 96-well plates were inoculated with HEK293T cells and incubated for 24 h. For the transient transfection of the cells GeneJuice transfection reagent (Novagen) was used according to the manufacturer's guidelines. The transfected expression cassettes were cloned into modified pEF1 vectors (Drerup et al. 2013). Each well was transfected with 110 ng of plasmid mixture (50 ng pEF1-RBOH; 30 ng of each effector; empty pEF1 vector to ensure equal loading). Transfected cells were incubated for 48 h and subsequently measured in a buffer consisting of Hank's Balanced Salt Solution (Gibco) with 62 µM L-012 and 60 µg / mL HRP. To stimulate SGN1, calyculin A, a phosphatase inhibitor, was added at the final concentration of 0.1 µM directly before the start of the measurements. ROS production was measured in a LB 943 Mithras$^2$ (Berthold) microplate reader and is presented as relative luminescence units $s^{-1}$ (RLU $s^{-1}$) in graphs. For all experiments, the values were gained from 6 wells measured in parallel and average values were plotted with S.D. Experiments have been repeated three times.

**Sample preparation for RNA-seq experiments**

Wild type, *sgn3-3* and *cif1-2 cif2-2* were grown on solid half-MS medium with mesh for 5 days and transferred onto fresh half MS medium with or without 100 nM CIF2. After 30, 120, or 480 min incubation, aerial parts were cut off and whole roots were collected. Samples were immediately frozen in liquid nitrogen and RNA was extracted using a Trizol-adapted Reliaprep RNA extraction Kit (Promega).

**RNA-seq library preparation and sequencing**

RNA quality was assessed on a Fragment Analyzer (Advanced Analytical Technologies, Inc., Ankeny, IA, USA). RNA-seq libraries were prepared using 1000 ng of total RNA and the Illumina TruSeq Stranded mRNA reagents (Illumina; San Diego, California, USA) on a Sciclone liquid handling robot (PerkinElmer; Waltham, Massachusetts, USA) using a PerkinElmer-developed automated script. Cluster generation was performed with the resulting libraries using the Illumina TruSeq SR Cluster Kit v4 reagents and sequenced on the Illumina HiSeq 2500 using TruSeq SBS Kit v4 reagents. Sequencing data were processed using the Illumina Pipeline Software version 2.20.

**RNA-seq data processing and analysis**

Purity-filtered reads were adapters and quality trimmed with Cutadapt (v. 1.8) (*51*). Reads matching to ribosomal RNA sequences were removed with fastq_screen (v. 0.11.1). Remaining reads were further filtered for low complexity with reaper (v. 15-065) (*52*). Reads were aligned against Arabidopsis thaliana TAIR10 genome using STAR (v. 2.5.3a) (*53*). The number of read counts per gene locus was summarized with htseq-count (v. 0.9.1) (*54*) using A. thaliana TAIR10 Ensembl 39 gene annotation. Quality of the RNA-seq data alignment was assessed using RSeQC (v. 2.3.7) (*55*).

Statistical analysis was performed for genes in R (R version 3.4.3). Genes with low counts were filtered out according to the rule of 1 count per million (cpm) in at least 1 sample. Library sizes were scaled using TMM normalization and log-transformed into counts per million or CPM (EdgeR package version 3.20.8) (*56*). PCA was computed using normalized values corrected for batch effect using limma function removeBatchEffect.

Differential expression was computed with limma-trend approach (*57*) by fitting all samples into one linear model. The batch factor was added to model matrix.

- Pairwise comparisons treated vs untreated per time point were assessed using moderated t-tests. The adjusted p-value is computed by the Benjamini-Hochberg method, controlling for false discovery rate (FDR or adj.P.Val).
- Differential expression of untreated mutant vs wt per time point was assessed using moderated F-test and Post-Hoc classification. The adjusted p-value is computed by the Benjamini-Hochberg method, controlling for false discovery rate (FDR or adj.P.Val).

- Differential expression of treated vs untreated over time paired by genotype was assessed using moderated F-test and Post-Hoc classification. The adjusted p-value is computed by the Benjamini-Hochberg method, controlling for false discovery rate (FDR or adj.P.Val).
- Interaction between time and treatment (paired by genotype, excluding SGN3 of the model) was assessed using moderated F-test. The adjusted p-value is computed by the Benjamini-Hochberg method, controlling for false discovery rate (FDR or adj.P.Val).
- Time effect in untreated conditions (paired by genotype, excluding SGN3 of the model) was assessed using moderated F-test. The adjusted p-value is computed by the Benjamini-Hochberg method, controlling for false discovery rate (FDR or adj.P.Val).

Genes were considered significant in further analysis if the adjusted p-value was equal or below 0.05 and the log2-fold change was >= 1. Heatmaps were constructed using the package ComplexHeatmap (v1.99.4, pearson distance for row clustering) (*58*). Genes were clustered using kmeans (factoextra v1.0.5, https://github.com/kassambara/factoextra) using a non-supervised approach resulting in 3 suggested clusters. We found that 4 inferred clusters split one gene cluster in a more sensible way and adjusted the number of clusters accordingly to four. GO analysis were conducted using the package topGO (v. 2.34.0, weight01 algorithm) (*59*). GO annotations were obtained through Biomart (version 2.40.0, Ensembl Plants release 43 - April 2019) (*60*). Genes for pathway specific heatmaps were obtained from the corresponding GO term through the Ensembl Plant database.

#### qPCR analysis

For the CIF2 peptide treatment, 5-days-old seedlings grown on half MS were moved to fresh 1/2 MS medium in the absence or presence of 100 nM CIF2 and incubated for 30 or 120 min. Otherwise, the plants were grown with mesh. Only root parts (around 100 mg) were collected at each time point and total RNA was extracted using a Trizol-adapted ReliaPrep RNA Tissue Miniprep Kit (Promega). Reverse transcription was carried out with PrimeScript RT Master Mix (Takara). All steps were done as indicated in manufactures' protocols. The qPCR reaction was performed on a Applied Biosystems QuantStudio3 thermocycler using a MESA BLUE SYBR Green kit (Eurogentech). All transcripts are normalized to *Clathrin adaptor complexes medium subunit family protein* (*AT4G24550*) expression. All primer sets are indicated in star methods.

#### Statistical analysis for experiments

All statistic-related analysis was done with R software(*61*)(https://www.r-project.org/). For multi-comparison analysis, one-way ANOVA was carried out and Tukey's test was subsequently performed.

**Fig. S1.**

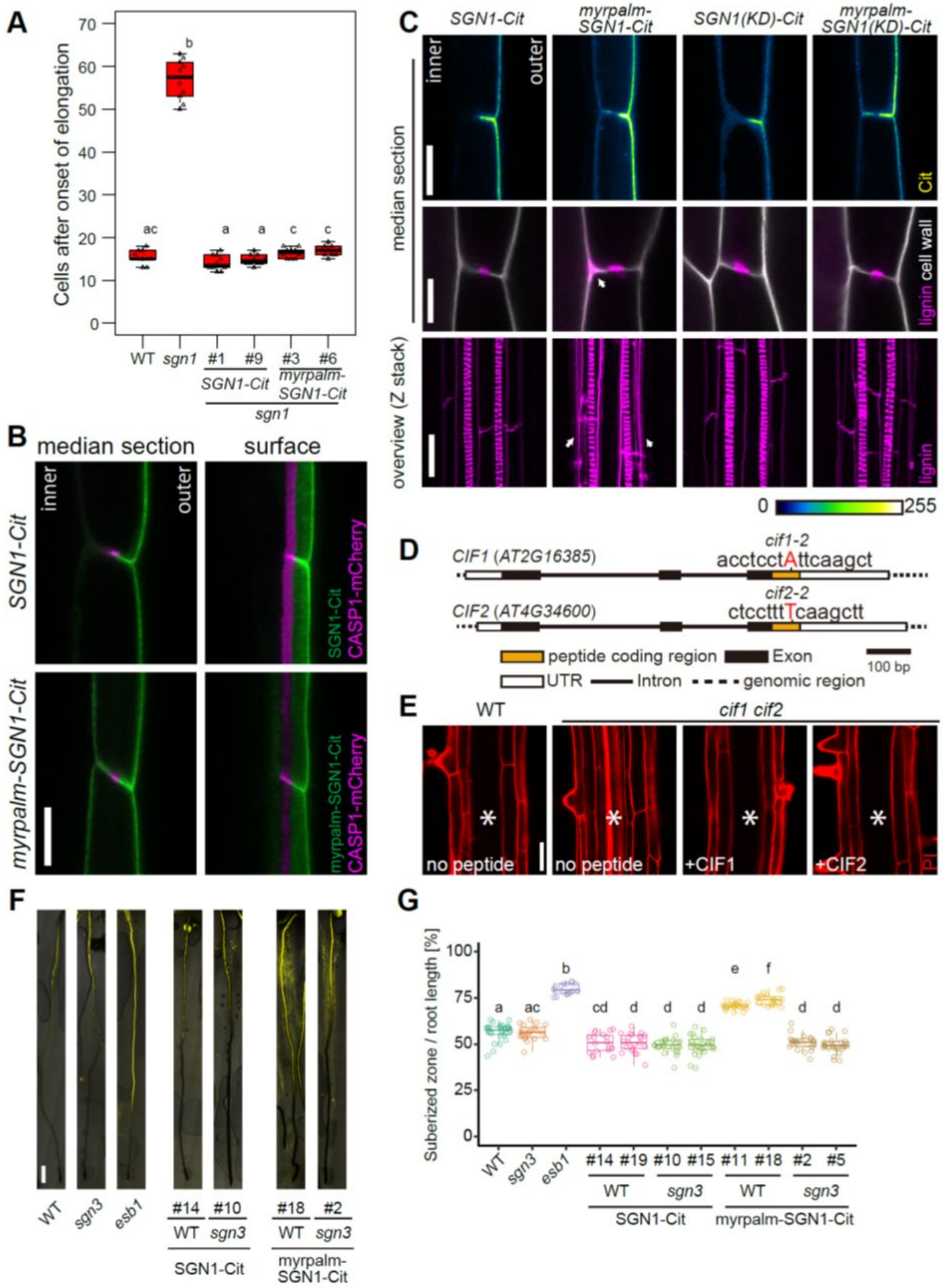

A
Cells after onset of elongation
WT sgn1 #1 #9 #3 #6
SGN1-Cit
myrpalm-SGN1-Cit
sgn1
B
median section
surface
inner
outer
SGN1-Cit
myrpalm-SGN1-Cit
SGN1-Cit CASP1-mCherry
myrpalm-SGN1-Cit CASP1-mCherry
C
SGN1-Cit
myrpalm-SGN1-Cit
SGN1(KD)-Cit
myrpalm-SGN1(KD)-Cit
median section
overview (Z stack)
Cit
lignin cell wall
lignin
D
CIF1 (AT2G16385)
cif1-2
acctcctAttcaagct
CIF2 (AT4G34600)
cif2-2
ctcctttTcaagctt
peptide coding region
Exon
100 bp
UTR
Intron
genomic region
E
WT
cif1 cif2
no peptide
+CIF1
+CIF2
PI
F
WT sgn3 esb1
#14 #10 #18 #2
WT sgn3 WT sgn3
SGN1-Cit
myrpalm-SGN1-Cit
G
Suberized zone / root length [%]
WT sgn3 esb1 #14 #19 #10 #15 #11 #18 #2 #5
SGN1-Cit
myrpalm-SGN1-Cit

**Fig. S1 Apolar SGN1 leads to ectopic lignin accumulation in endodermal cells**

(A) PI penetration assay. Scoring of number of cells after onset of cell elongation until PI signal is excluded from inner side of endodermal cells (n= 10). Different letters indicate significant statistical differences ($p < 0.01$, One-way ANOVA and Tukey test).

(B)Localization of CASP1-mCherry, driven by CASP1 promoter, in *pCASP1::SGN1-Cit* or *pCASP1::myrpalmSGN1-Cit* transgenic lines. Scale bar = 10 μm.

(C) Localization patterns of SGN1 (WT, kinase dead (KD)), myrpalm-SGN1 (WT and KD) and lignin deposition patterns in each indicated transgenic lines. Arrowhead indicate excess lignification. Scale bars are 10 μm in SGN1-Cit, 5 μm in lignin and calcofluor pictures, 20 μm in overview of lignin deposition.

(D) One-base pair insertion sites of *cif1-2* and *cif2-2*. Red letters indicate inserted base in each locus.

(E) PI penetration phenotype of the *cif1cif2* double mutant with or without 100 nM peptide treatment. Seedlings were germinated on the medium with or without peptides. Asterisk indicates the stele. Scale bar = 40 μm.

(F) Whole root views of suberin deposition patterns in polar- or apolar-SGN1 transgenic lines. *esb1* (*enhanced suberin 1*) is shown as a representative over-suberized mutant. Scale bar = 500 μm.

(G) Quantification of ratio of suberized zones and root lengths in each mutant or transgenic line. *esb1* is shown as a representative over-suberized mutant. Different letters are indicating statistically significant difference (n = 16 – 36 roots, $p < 0.01$, ANOVA and Tukey test).

**Fig. S2.**

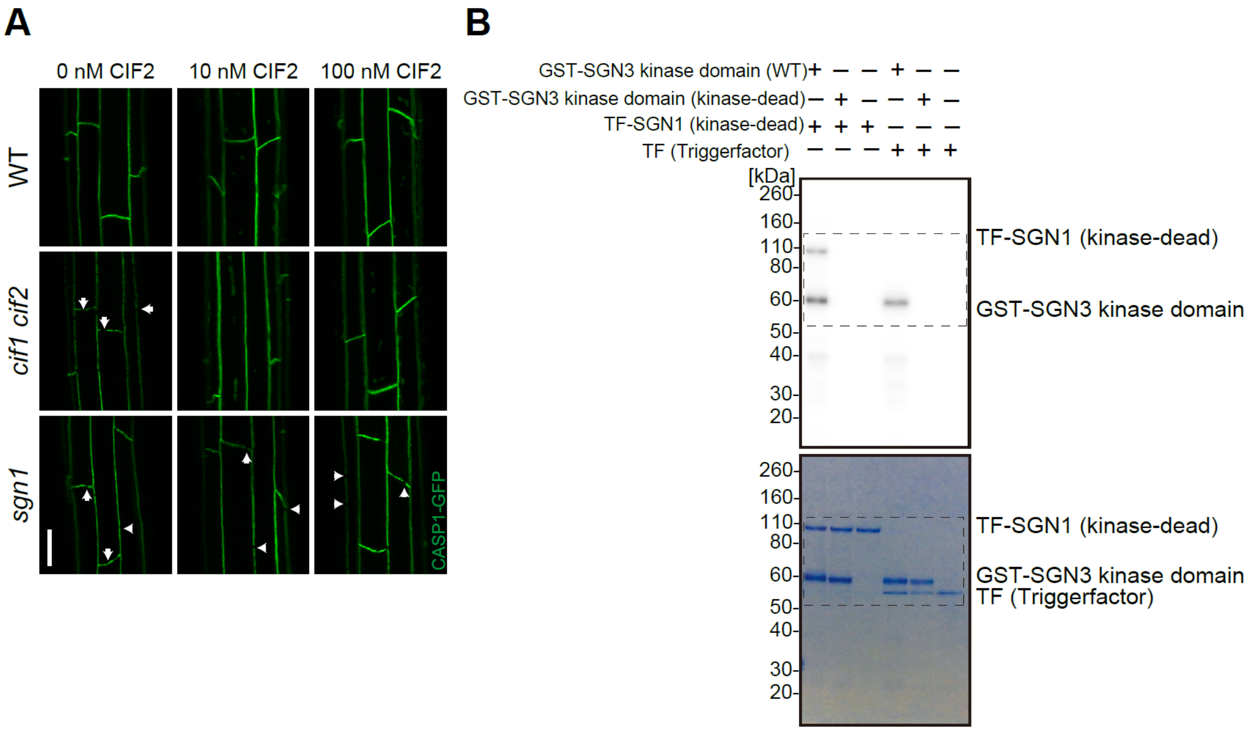


**Fig. S2 SGN1 acts as a transducer of CIF2 signaling and is phosphorylated by the SGN3 receptor**

(A) Representative images of CASP1-GFP localization pattern after 5-day in the presence or absence of 10 or 100 nM CIF2. Arrowheads highlight some of the discontinuities in the mutant CSDs. Scale bar = 20 $\mu$m.

(B) Full scan of the gels corresponding to Figure 2C. The regions in the dashed boxes are presented in Fig. 2C.

**Fig. S3.**

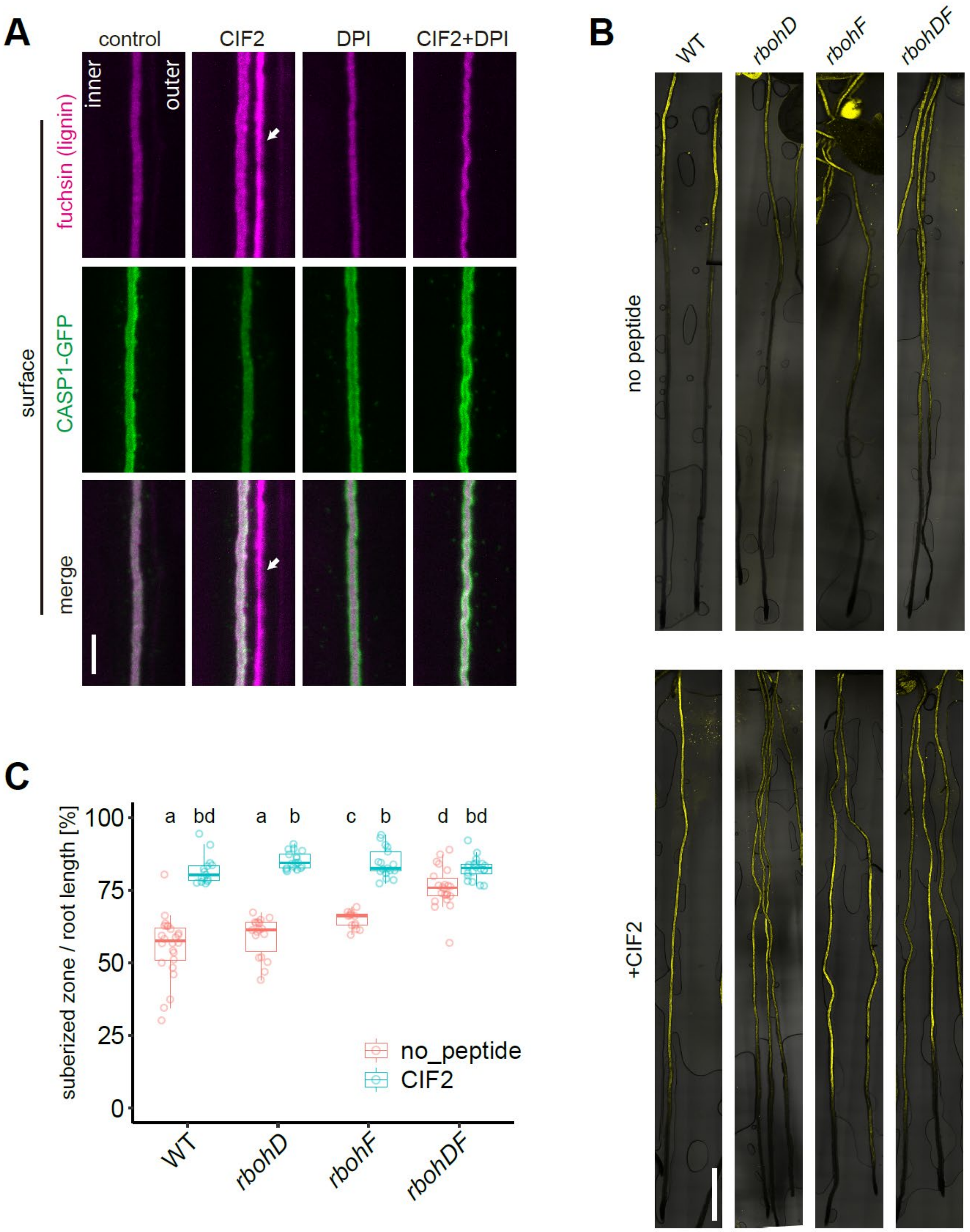
A
control
CIF2
DPI
CIF2+DPI
inner
outer
fuchsin (lignin)
CASP1-GFP
merge
surface
B
WT
rbohD
rbohF
rbohDF
no peptide
+CIF2
C
suberized zone / root length [%]
100
75
50
25
0
a
bd
a
b
c
b
d
bd
no_peptide
CIF2
WT
rbohD
rbohF
rbohDF

**Fig. S3: Both RBOHD and F are required for CIF2-induced excess lignin accumulation**

(A) Co-treatment experiments with the NADPH oxidase inhibitor DPI (Diphenyleneiodonium chloride) and CIF2. Pretreatment was done on medium with or without DPI for 30 min and seedlings were then transferred to each medium. The seedlings were incubated for 2 hours in each condition. Arrowheads indicate excess liginification on the cortex-facing side. Scale bar = 5 $\mu$m.

(B) Whole root views of suberin deposition patterns in the *rbohd* and *rbohf* mutants with or without CIF2 treatment. 5-day-old seedlings were treated for 24 hours with or without CIF2 and stained. Scale bar = 1 mm.

(C) Quantification of the ratio of suberized zones to root lengths in each mutant from (B). (n= 13-24 roots). Different letters indicate statistical significance (One-way ANOVA, Tukey's test).

**Fig. S4.**

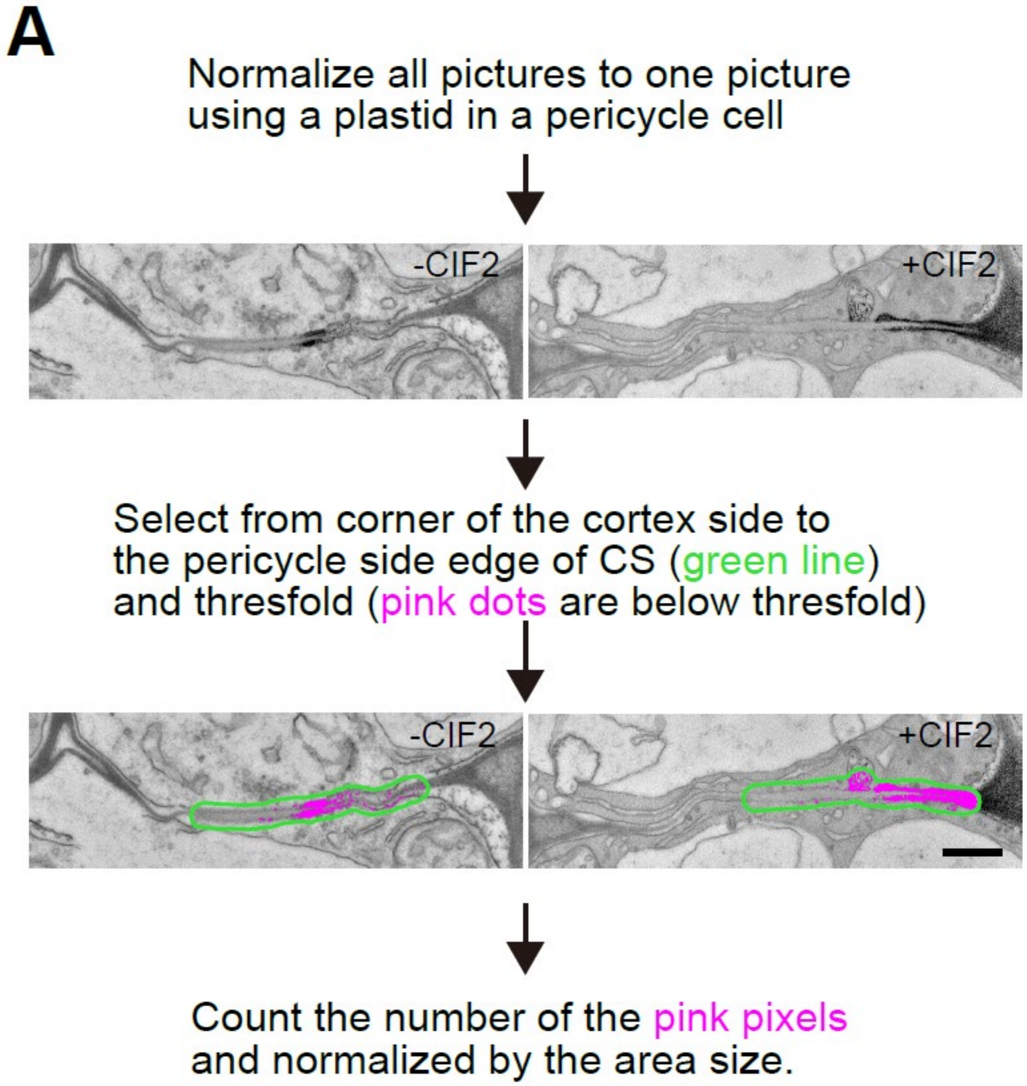


**Fig. S4: ROS production is enhanced by SGN3/CIFs and requires RBOHD and F**

(A) A schematic illustrating the protocol for pixel area quantification of the ROS production sites. Pictures were normalized to a picture of non-treated WT. Following the normalization, the area was chosen manually from cortex side corner to the end of the CS at the pericycle side. Pixels below threshold were marked as pink dots and counted. For more details, see the materials methods part. Scale bar = 500 nm.

**Fig. S5.**

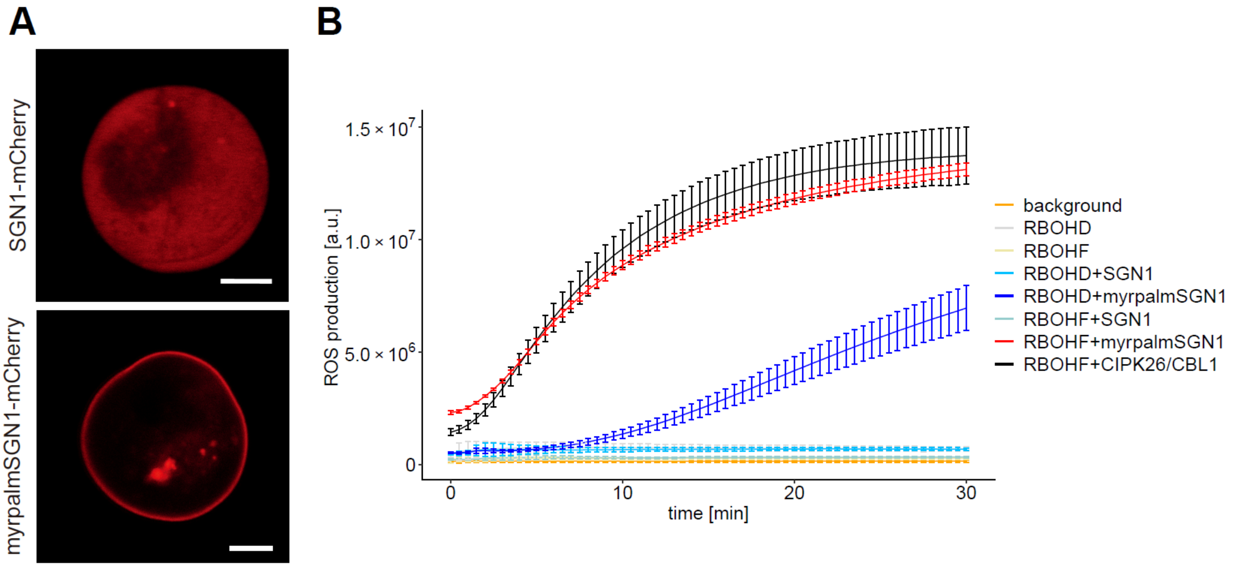


**Fig. S5 SGN1 directly activates NADPH oxidases in a cellular context**

(A) Localization patterns of SGN1-mCherry of myrpalm-SGN1 in HEK293T cells. Myrpalm-SGN1 is efficiently recruited to the plasma membrane, while wild-type SGN-mCherry fusions remain in the cytoplasm. Scale bar = 5 $\mu$m.

(B) Independent HEK cell ROS production assay. The phosphatase inhibitor CalyculinA was added directly before the start of the measurements. Each data point represents the mean of six wells analyzed in parallel, bars indicate S.D.

Fig. S6.

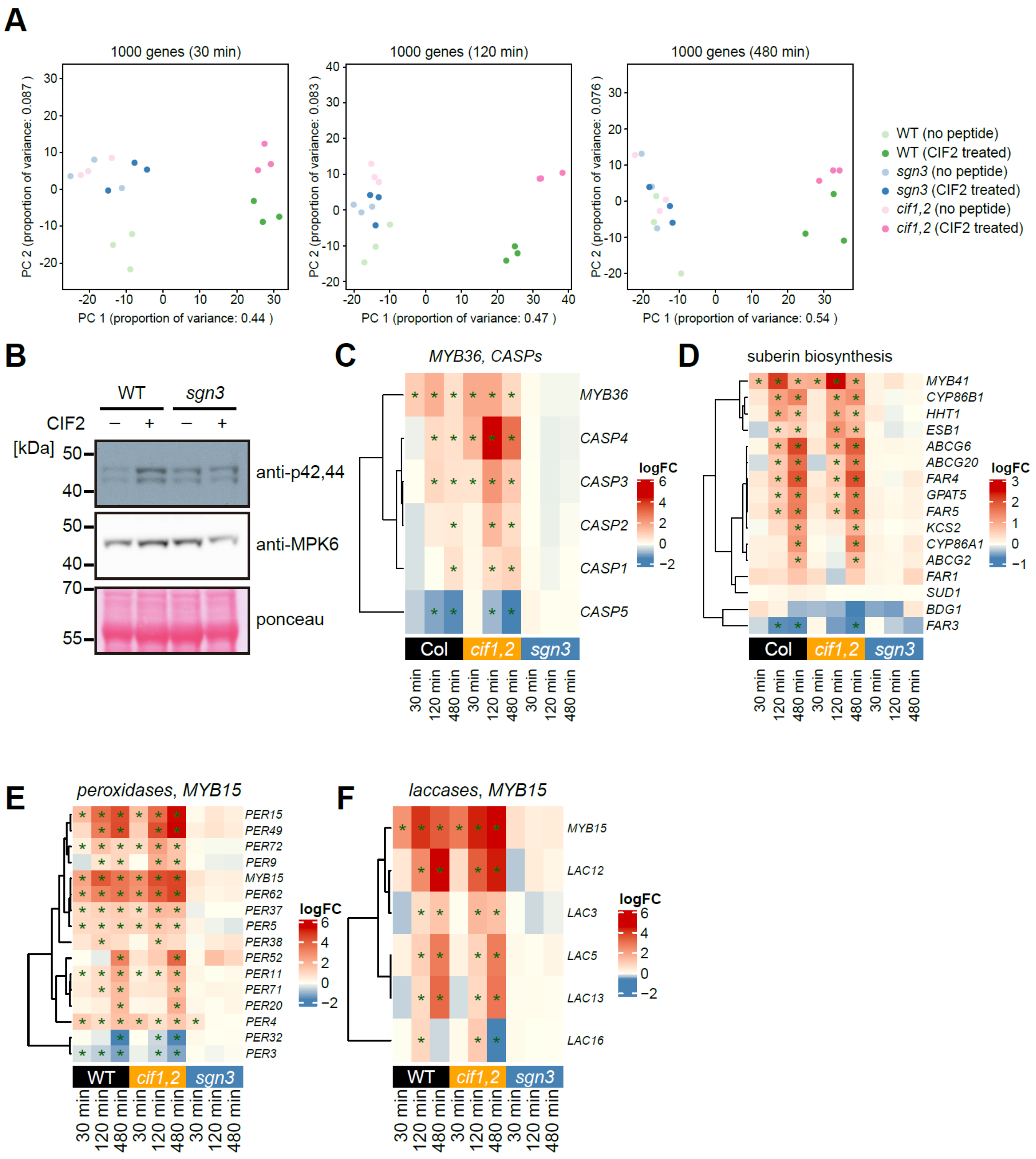


**Fig. S6 CIF2 induces large-scale transcriptional changes to remodel the cell walls**

(A) PCA analysis of most differentially expressed genes. The 1000 most differentially expressed genes are clustered by condition at each time point for all replicates.

(B) Immunoblot with phospho-specific antibody against p42,44. Seedlings were treated with or without 1 μM CIF2 peptide for 15 min. Ponceau S stained membranes and IB with anti-MPK6 antibody were shown as loading control. This experiment was repeated three times with independent biological samples with the same result.

(C) Heatmaps of *MYB36* and *CASPs* expression fold-changes with or without the peptide treatment at the indicated time points. Asterisks indicate significant differentially regulated transcripts ($p \leq 0.05$) at each condition.

(D) Heatmaps of *MYB41* and suberin biosynthesis-related gene expression fold changes with or without the peptide treatment at the indicated time points. Asterisks indicate significant differentially regulated transcripts ($p \leq 0.05$) at each condition.

(E) Heatmaps of *MYB15* and *PEROXIDASES* expression fold changes with or without the peptide treatment at the indicated time points. Asterisk indicates Asterisks indicate significant differentially regulated transcripts ($p \leq 0.05$) at each condition.

(F) Heatmaps of *MYB15* and *LACCASES* expression fold changes with or without the peptide treatment at the indicated time points. Asterisks indicate significant differentially regulated transcripts ($p \leq 0.05$) at each condition.

**Table S1.**

Differentially expressed genes from heatmap presented in Fig.6D

**Table S2.**
A list of enriched GO terms of clusters presented in Fig.6D

**Movie S1.**
Time lapse movie of CIF2 treatments with or without cycloheximide (CHX) on CASP1-GFP in *cif1 cif2*.
